# Temperature-related biodiversity change across temperate marine and terrestrial systems

**DOI:** 10.1101/841833

**Authors:** Laura H. Antão, Amanda E. Bates, Shane A. Blowes, Conor Waldock, Sarah R. Supp, Anne E. Magurran, Maria Dornelas, Aafke M. Schipper

**Affiliations:** Research Centre for Ecological Change, Organismal and Evolutionary Biology Research Programme, University of Helsinki, Helsinki, Finland; Centre for Biological Diversity, University of St Andrews, St Andrews, Scotland; Department of Ocean Sciences, Memorial University of Newfoundland, Newfoundland, Canada; German Centre for Integrative Biodiversity Research (iDiv), Halle-Jena-Leipzig, Germany; Ocean and Earth Science, National Oceanography Centre, University of Southampton, Southampton, UK; Department of Life Sciences, Natural History Museum, London, UK; Data Analytics Program, Denison University, Granville, OH, USA; PBL Netherlands Environmental Assessment Agency, The Hague, The Netherlands; Department of Environmental Science, Institute for Water and Wetland Research, Radboud University, Nijmegen, The Netherlands

**Keywords:** temperature change, species richness, abundance, species gains, species losses, biodiversity change, climate change

## Abstract

Climate change is reshaping global biodiversity as species respond to changing temperatures. However, the net effects of climate-driven species redistribution on local assemblage diversity remain unknown. Here, we relate trends in species richness and abundance from 21,500 terrestrial and marine assemblage time series across temperate regions (23.5-60.0°) to changes in air or sea surface temperature. We find a strong coupling between biodiversity and temperature changes in the marine realm, which is conditional on the baseline climate. We detect increases in species richness with increasing temperature that is twice as pronounced in warmer locations, while abundance declines with warming in the warmest marine locations. In contrast, we did not detect systematic temperature-related richness or abundance trends on land, despite a greater magnitude of warming. We also found no evidence for an interaction between biodiversity change and latitude, further emphasizing the importance of baseline climate in structuring assemblages. As the world is committed to further warming, significant challenges remain in maintaining local biodiversity amongst the non-uniform inflow and outflow of “climate migrants” across distinct regions, especially in the ocean.

---

Climate change is driving a reorganization of ecological communities as species track changes in air and ocean temperatures. However, global warming is not unfolding evenly across the planet, and this heterogeneity is layered over the uneven distribution of biodiversity. Species benefiting from warming exhibit abundance increases and expand their geographic ranges^1–5^. Conversely, species that are more thermally restricted may decline with warming, as individuals die or move to more suitable locations^4,6,7^. The latter is typically expected for tropical species, since they have narrower thermal tolerances than temperate species, and live closer to their upper thermal limits^6,8–11^. In contrast, mid-to high-latitudes undergoing warming may provide suitable habitat for species expanding their ranges poleward^4,12^. As the tropics hold the majority of the world’s species^13^, lower-latitude temperate regions undergoing warming may experience larger increases in species richness and abundance compared to temperate locations at higher latitudes (Fig. 1). Biodiversity change may further unfold differently depending on the baseline climate, i.e. the effects of warming might differ between initially warmer *versus* colder regions^3,6,12,14^. For instance, warm temperate regions may offer the first habitats suitable for climate immigrants.

**Figure 1.**
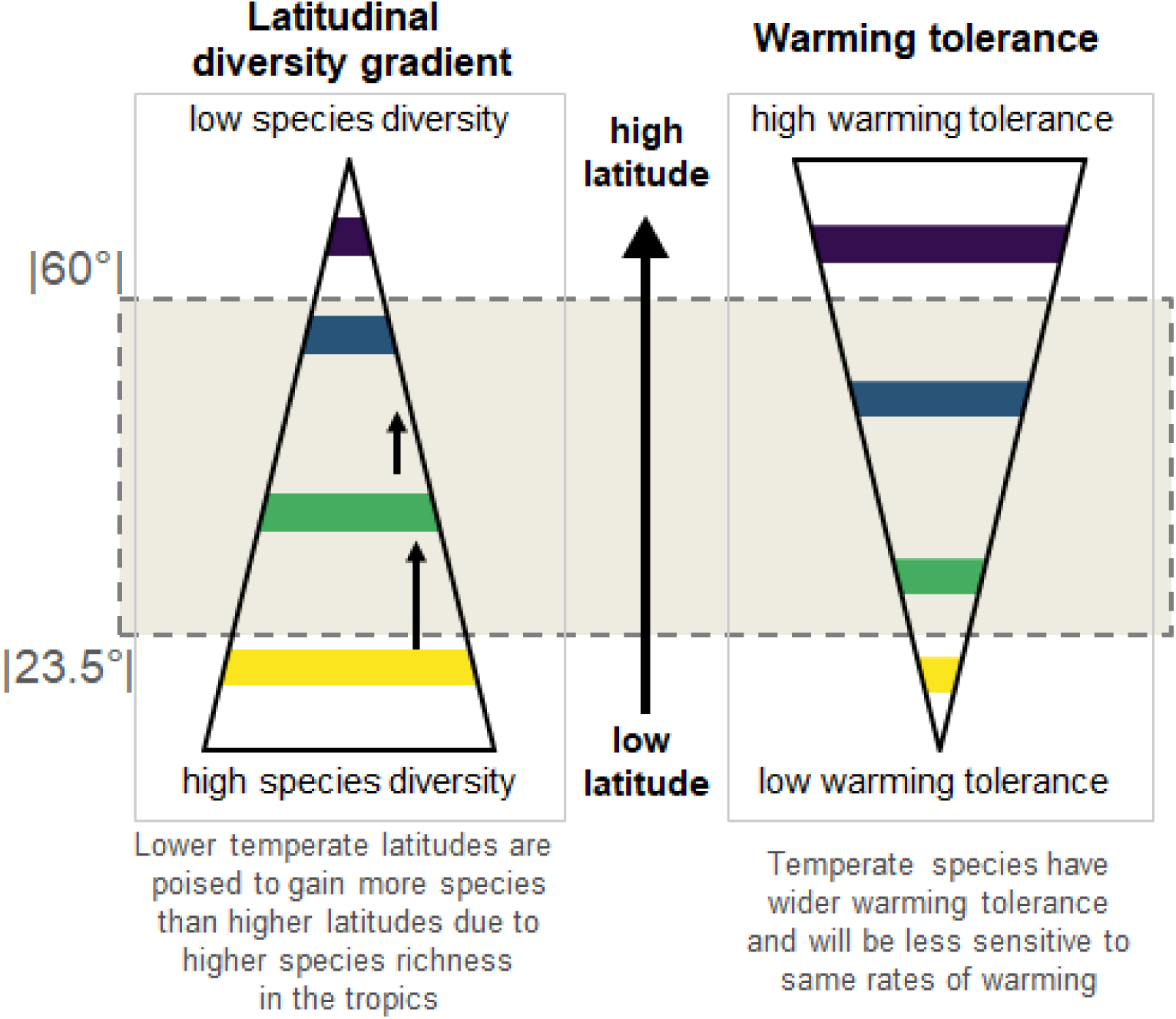
Mechanisms underpinning the expectation that temperature-related biodiversity change unfolds unevenly across the planet. (within the temperate latitudinal band where our data fall), stemming from the latitudinal gradients in species diversity (decrease with latitude) and species warming tolerances (or thermal safety margins (TSM); increase with latitude).

Warming-induced biodiversity change may also be relatively strong in the ocean^3,15,16^. Marine species are highly responsive to temperature change and can track changing isotherms with fewer barriers to dispersal, compared to terrestrial species^3,14–19^. Moreover, the availability of thermal microrefugia is limited in the ocean, while in terrestrial ecosystems organisms can seek shade or burrow in soil to buffer the effects of warming^17,20^. Therefore, the effects of temperature change are expected to be more immediate and directly detected for marine ecosystems, an expectation which is supported by a growing literature which has quantitatively compared the responses of individual species between marine and terrestrial realms^3,14–16^. However, the net effects of temperature-related species’ movements and abundance changes on assemblage-level diversity have not yet been systematically quantified across realms.

Here, we quantify temperature-related species richness and total abundance change in marine and terrestrial assemblages across temperate regions of the planet (23.5°-60.0° absolute latitude). Specifically, we test two predictions for the effects of temperature change on assemblage-level diversity: (1) species richness and total abundance will increase with warming, and such increases will be greatest across relatively warm regions that border the species-rich tropics; and (2) the coupling of assemblage and temperature change will be tighter in the ocean than on land. We focus on assemblage-level trends rather than species-specific responses, and quantify changes in both total abundance and species richness. These two metrics can be decoupled from each other, and abundance is typically more responsive than richness^21,22^. We further disentangle richness change into species gains and losses to better understand the underlying dynamics of temperature-related biodiversity change. To test our expectations, we used the largest database of assemblage time series, BioTIME^23^, which includes studies for plants, invertebrates, birds, mammals, and fish, to quantify biodiversity trends. These assemblages consist of co-occurring species systematically sampled through time. Since spatial extent varied among studies in BioTIME, we harmonized the biodiversity observations to a common spatial resolution to minimise the influence of variation in spatial extent on our results^24^; this allowed us to quantify the effect of temperature change at a standardised resolution across regions and realms. We first estimated trends in biodiversity and temperature separately, and then quantified the relationships between the two.

Specifically, for each study we allocated individual samples to 96 km^2^ hexagonal grid cells based on their location (Methods; ^24^). For studies with large extents, we created new equal-extent assemblage time series by allocating the samples to different grid-cell by study combinations (thus keeping the integrity of each sample and study within each grid cell). We used these new spatially harmonized assemblage time series in our analysis, selecting data from temperate regions only (since these are the better sampled regions within BioTIME). We then selected time series with at least five years of sampling (mean=9.2 years), yielding 21,500 assemblage time series across both realms (19,875 marine and 1,625 terrestrial from 156 original studies; Fig. S1, Table S1). Because the number of samples can vary from year to year within each time series, we used sample-based rarefaction to equalise sampling effort among years and then quantified trends in richness, total abundance, and number of species gains and losses. For the same locations and for the time spans matching the duration of the biodiversity monitoring periods, we extracted mean monthly temperature records from HadCRUT4^25,26^ and estimated the corresponding rates of sea surface or air temperature change per year. We then quantified the relationships between changes in biodiversity and changes in temperature with meta-analytical Bayesian hierarchical models, allowing responses to vary among taxonomic groups. To test our expectations, we included an interaction term between temperature change and latitude or long-term average temperature (i.e. baseline climate) in our models, fitted separately for the marine and terrestrial realms.

## Results

Temperature trends were highly variable, with locations at similar latitudes exhibiting different directions and magnitudes of change (Fig. 2a). Yet, both sea surface and air temperatures increased on average at the locations and time periods of our study, even though the majority of our time series spanned less than 10 years. The warming signal was more pronounced on land than in the ocean (Fig. 2b; average mean temperature change rate was 0.022 °C/year on land, *versus* 0.012 °C/year in the ocean).

**Figure 2.**
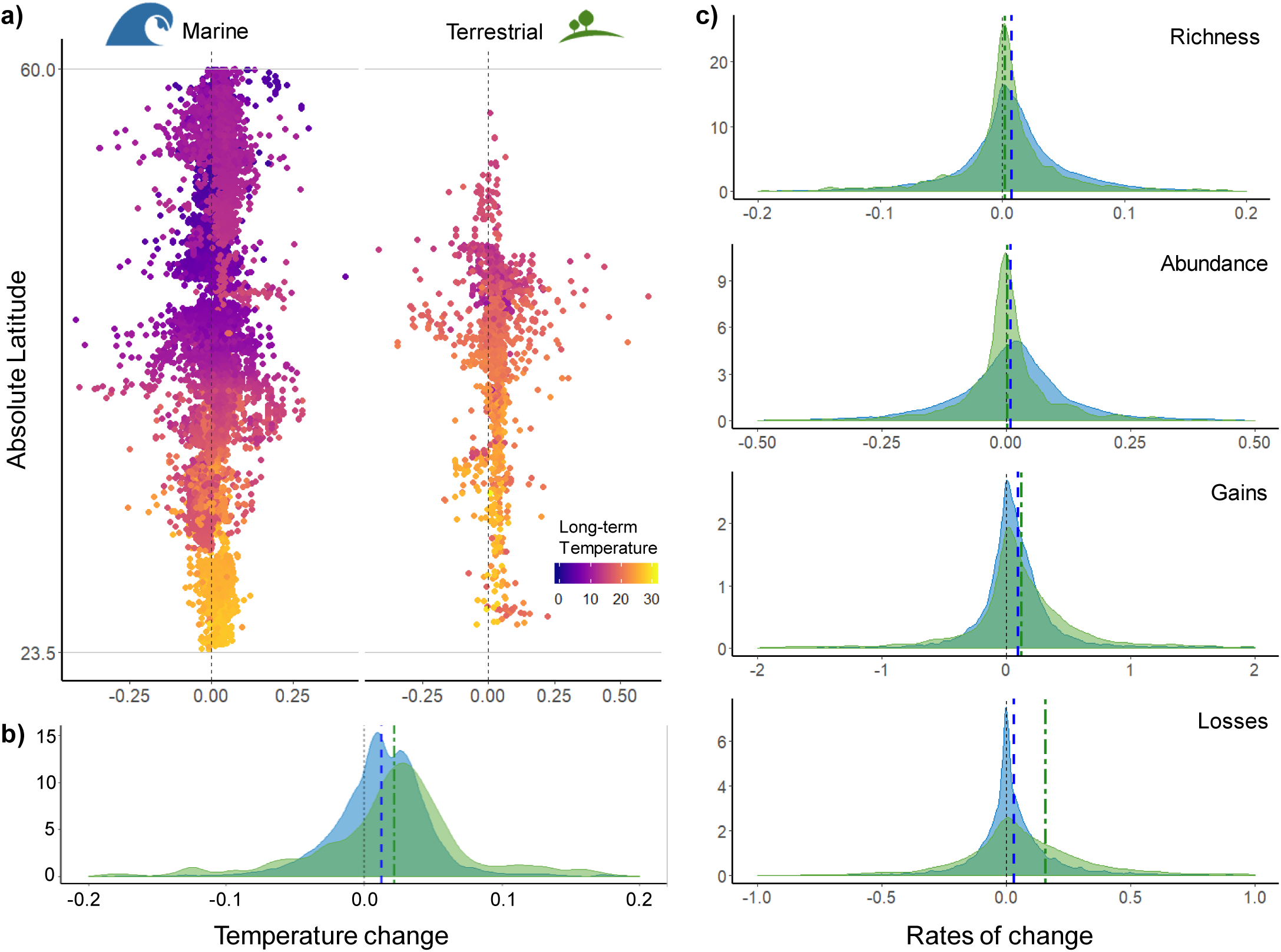
Variation in temperature and local biodiversity trends across the time series. (marine in blue, n= 19,875; terrestrial in green, n= 1,625). **(a)** Each point indicates the rate of temperature change (°C/year) for a specific time series, coloured according to the long-term average temperature. There was no clear latitudinal pattern in temperature change: while the majority of locations in both realms experienced warming, and more so for terrestrial locations **(b)**, many locations experienced cooling during the period examined. Local biodiversity change estimates (rate/year) also exhibited wide variability **(c)** (note the different scales for the different metrics; x-axes were truncated to improve clarity). Tick dashed vertical lines indicate the overall mean per realm in all the density plots. The biodiversity time series locations cover numerous habitats and biomes (Fig. S1), and sample a large range of the planet’s long-term average temperature gradient.

Biodiversity change was also highly variable among the assemblage time series (Fig. 2c). Yet, despite the variability in both temperature and biodiversity trends, coherent macroecological signals emerged, but only in the ocean and conditional on the baseline climate (Fig. 3, Table S2). In the marine realm, we found an overall increase in species richness with warming that was twice as pronounced in warmer locations, underpinned by higher rates of species gains and lack of systematic changes in species losses (Figs. 3, 4 and S2). Additionally, warming coincided with losses of individuals in the warmest marine locations, whereas cooler locations tended to gain individuals with increasing temperature (Figs. 3, 4 and S2). In contrast, no systematic biodiversity responses emerged on land, where the 95% credible intervals overlapped zero for all the biodiversity metrics included (Fig. 3, Table S2). Our analysis clearly highlights the fundamental role of climate baselines in modulating biodiversity responses (particularly in the ocean), given that latitude showed no or very weak interacting effects with temperature change (Fig. S3).

**Figure 3.**
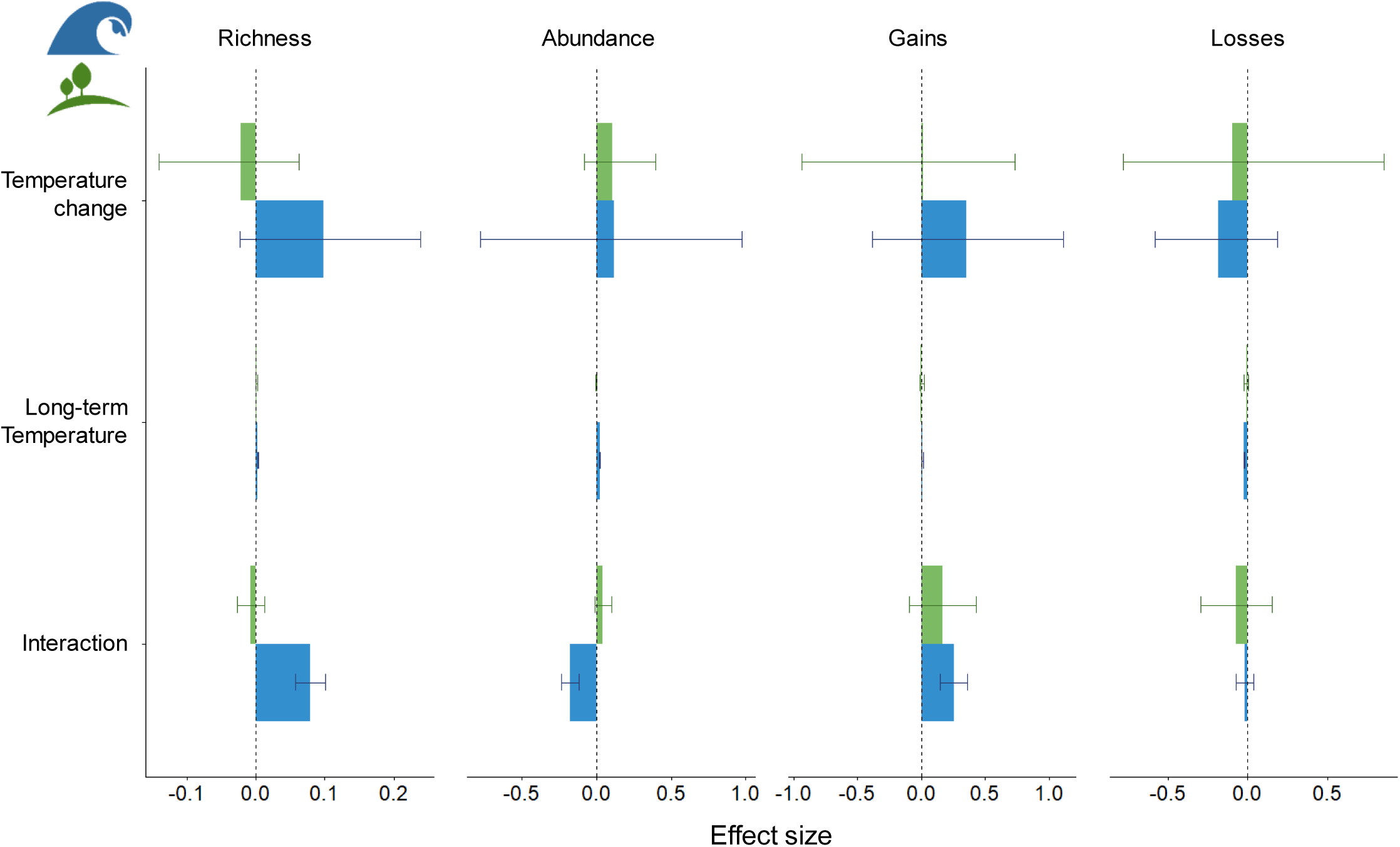
Biodiversity responses to the interacting effects of temperature change and long-term average temperature (i.e. baseline climate). Marine locations (blue) exhibited stronger responses compared to terrestrial locations (green), while baseline climate modulated these responses in divergent directions. Bars represent the effect sizes and whiskers indicate the 95% credible intervals estimated from the Bayesian meta-analysis (note the different scales for the different metrics); estimated parameters were considered to represent signals in the responses when the credible intervals did not include zero (Table S2).

**Figure 4.**
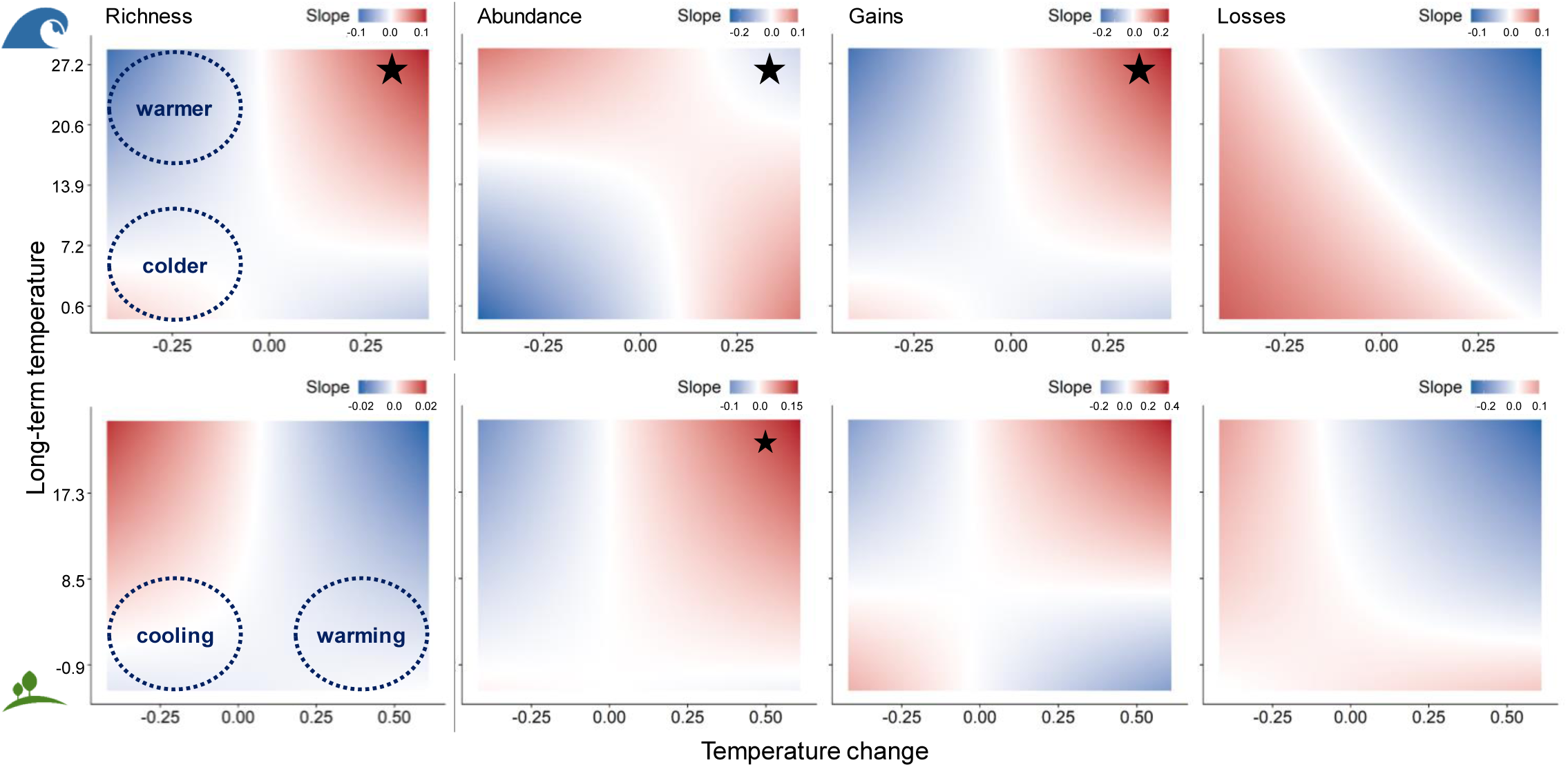
Biodiversity responses to temperature change across different baseline climates. Each panel depicts the rate and direction of biodiversity change depending on the temperature change experienced (cooling *versus* warming) and on the long-term average temperature (colder *versus* warmer), where red indicates positive slopes for the biodiversity response, and blue indicates negative slopes (note the different scales across the different metrics), for marine (top row) and terrestrial realms (bottom row). Stars indicate when the 95% credible intervals for the interaction term from the meta-analytical models did not overlap zero (Fig. 3, Table S2; smaller star for abundance change on land indicates 95% CI marginally overlapping zero).

The different responses between the two realms were robust to uneven sampling in terms of number of locations and latitudinal range (Fig. S4). Our results were also consistent across different temperature variables (long-term annual and maximum temperature, and annual mean temperature of the first year) and between different temperature databases for quantifying climate baselines (Fig. S3; see Methods).

## Discussion

We reveal striking differences in warming-related biodiversity change between marine and terrestrial realms, with a much stronger signature of warming on marine assemblages. Our results are unlikely due to confounding factors, given that climate change is poorly spatially correlated with other anthropogenic drivers of change for both marine and terrestrial realms^27^. Further, temperature is expected to be a strong driver of biodiversity change^28–30^.

The warming-related increase in marine species richness is consistent with the expectation that as the Earth’s climate warms, temperate regions undergoing warming will receive an influx of species tracking suitable temperatures, and increases in warm-affinity generalists^2,7,8,18,28^. This warming-related increase in species richness is likely, in part, underpinned by species from lower latitudes shifting their ranges poleward to avoid exceeding their upper thermal limits^4,12^. Indeed, projected rates of extirpation in response to recent and future warming are highest for tropical marine species^8,31^. Here, we found that species gains outpaced losses in the warmest locations in the ocean where temperature has also increased. This finding is consistent with asymmetrical responses between species range edges, with faster colonisations expected following climatic niche expansion, and with slower local extinctions linked to delayed responses at species trailing edges^3,4^. The prevailing influx of species with warming is likely to lead to novel biotic communities and interactions as species distributions are re-shuffled, with potentially far-reaching consequences for ecosystems functioning^3,8,31,32^.

Our results further highlight that substantial loss of individuals can occur simultaneously with increases in richness, illustrating that temperature-related changes in richness and abundance can be decoupled. Abundance declines may be occurring for more thermally restricted species, owing to reduced performance and population declines, as critical thermal thresholds are crossed^4,6,7^, for instance related to the adverse effects of increasing temperatures on metabolic rates and primary production^28,33^. Influxes of climate immigrants can also drive local declines in populations due to greater grazing and predation rates, e.g.^34,35^. The abundance declines across our warmest locations in the ocean suggest warming-related destabilization of populations possibly reflecting reductions in the carrying capacity of marine ecosystems.

We also find strong modulating effects of baseline climatic conditions on both abundance and richness responses that are not attributable to latitude *per se*. These findings highlight that rising temperatures in locations that are already warm in comparison to other regions from similar latitudes can lead to greater assemblage restructuring. This may reflect the patchiness in temperature regimes across similar latitudes, for example due to altitudinal or depth gradients, proximity to the coast, or ocean currents. The baseline climate therefore emerges as a major predictor of temperature-related biodiversity change in marine systems.

Overall, and despite faster warming on land, terrestrial assemblages did not show systematic responses in richness or abundance with temperature change. The stronger responses observed for marine assemblages are consistent with reported faster range shifts in the ocean and higher sensitivity of marine organisms to temperature change compared to terrestrial species^3,12,14–16^. Our findings are also consistent with warming-related local extirpations being twice as common in the ocean as on land^15^. The lack of systematic assemblage-level change associated with warming on land might be due to greater thermoregulation capacity and wider thermal safety margins of terrestrial taxa^9,15,16^. Additionally, compared to seascapes, higher landscape complexity enables terrestrial species to exploit thermal microhabitats, thus allowing for the persistence of local populations for longer periods^9,20^. Indeed, access to thermal refugia was reported to be a fundamental factor underlying the relatively low vulnerability of terrestrial ectotherm species to warming, and emphasizes the potential deleterious combined effects of warming and land-use changes^15^. Finally, a weaker link between assemblage responses and temperature change on land may be due to other factors, such as land-use change and moisture availability, posing stronger constraints on local biodiversity. Nonetheless, the smaller magnitude or slower responses of terrestrial species to temperature change^5,12,15,24,36,37^, combined with the faster rates of warming on land, indicate a potentially higher risk of climatic debt (i.e. response lags) among terrestrial *versus* marine taxa^3,12,14,19,36,38–40^. Additional research with higher-resolution temperature data matching the scale of organisms’ responses is needed to better quantify terrestrial assemblages responses to temperature change, and these developments remain a major challenge for many different taxa.

Our analyses provide a first step towards explaining divergent patterns of assemblage-level biodiversity change across the planet^24,41^. Overall, our results provide strong support the prediction that divergent biodiversity trajectories across latitude may arise as a consequence of global warming, with polar and temperate regions likely acting as “sinks”, and tropical regions as “sources”^6,8,31^. In turn, these responses could prompt a shift in the latitudinal diversity gradient towards higher latitudes, with faster rates of change in the ocean. While we focused here on temperate regions, tropical and polar biomes are predicted to undergo severe restructuring in response to temperature change, albeit along different trajectories^6,8,12^. However, lack of sufficient biodiversity monitoring data for tropical and polar systems^23,42^ hampers a comprehensive assessment of assemblage-level responses to temperature change in these regions.

Future global warming impacts on biodiversity are likely to exceed and potentially diverge from the changes revealed here^8,31,33,43–45^. Indeed, initial increases in richness and abundance in response to warming may be followed by losses if warming continues^7,46,47^. These declines in marine systems may affect food security and livelihoods of human populations that depend on the ocean^7,48^. Additionally, while a consistent signal of temperature change was not evident on land, future impacts on terrestrial assemblages are expected from continuing rising temperatures, extreme heat events, fires, and lack of moisture^33,44,49^. Because the Earth is committed to further warming, a systematic reduction of greenhouse gas emissions alongside efforts to further prevent habitat loss and improve habitat connectivity will be fundamental to allow species to track suitable climates across increasingly impacted land- and seascapes and to avoid severe biodiversity disruption and loss.

## Methods

### Biodiversity data and trends

BioTIME^23^ is currently the largest global database of assemblage time series, including 386 individual studies (Study ID; plus extended data sources) across different taxonomic groups, holding over 12 million records of abundance for over 45 thousand species. For this analysis, we only included studies reporting counts of individuals per species in terrestrial and marine systems. We excluded freshwater studies as these are too few to confidently analyse biodiversity trends across taxa and different regions.

Each study is comprised of distinct samples (i.e. individual plots, transects, tows, etc. sampled at a given time), and the number of samples can vary among years within each study. As the spatial extent varies among studies, we gridded those studies that had large extents and multiple sampling locations into hexagonal cells of ~96km^2^; many studies were not partitioned because they were contained within a single cell^24^. Specifically, each sample was assigned to different combinations of study ID x grid cell based on their latitude and longitude, resulting in new assemblage time series (each with multiple samples across years). These new time series were given a unique identifier that was the concatenation of the study ID and the grid cell reference number, and thus contained samples from only one study ‒ i.e. the integrity of each study and each sample were maintained. This process allowed us to relate biodiversity and temperature trends at a standardized resolution. To minimise the effect of unobserved species on estimates of biodiversity change, we calculated the abundance-based coverage^50^ of each annual sample within each time series, and removed all samples with coverage less than 0.85. To be able to estimate reliable biodiversity trends, we restricted our analysis to time series sampled in at least five years (not necessarily consecutive). Because the number of samples can vary among years, we used sample-based rarefaction^51^ to standardise the number of samples among years for each time series before calculating the biodiversity metrics. Specifically, we identified the minimum number of samples taken in each year within each assemblage time series; this minimum was then used to randomly sample each year down to that number of samples. Finally, given the paucity of data representing polar and tropical regions, we excluded these regions (based on absolute latitudinal cut-offs at 60° and 23.5°, respectively). This process yielded 21,500 assemblage time series representing 156 original studies (Table S1) between 1900 and 2016, across 19,875 marine and 1,625 terrestrial locations. The average number of years sampled across the time series was 9.2 years, with the longest time series spanning 97 years (Fig. S5).

To quantify rates of biodiversity change, we calculated linear trends over time for species richness (logS), total abundance (logN), number of species gains and species losses. Counts of gains and losses retained species identity information, and were quantified based on comparison with the first year sampled in each time series. For losses, a positive slope means the number of species lost from a location is increasing through time; negative slopes represent time series where the magnitude in species losses decreased over time. We repeated the sample-based rarefaction process described above 199 times for each time series, recorded the values and took the median for each biodiversity metric in each year, in order to reduce the effect of any outlier samples on our estimates. We used ordinary least squares regression because we were interested in the long-term direction and magnitude of the biodiversity trends, and to allow us to compare the rates of change among locations, realms and metrics. We retained the estimated slope and standard error for each time series for use in our second-stage meta-analytic models.

### Temperature data and trends

We focus on temperature as a climate variable because of its influence on every level of biological organization, from individual metabolic rates to ecological communities’ dynamics and structure^28–30^. We extracted temperature records from HadCRUT4^25,26^, specifically the HadSST3 data for marine Sea Surface Temperature (SST) on a 1° resolution, and the CRUTEM4 data for air temperature on land on a 0.5° resolution. We did not harmonize the spatial resolution between the two data sources because we wanted to use the best available data in each realm. For the location of each biodiversity time series, we extracted monthly mean temperature records for the duration of the biodiversity monitoring period (Year_start_:Year_end_), and estimated mean temperature trends using generalized additive models (GAM), including a temporally autocorrelated error structure (package mgcv^52^). This also allowed us to assess if accounting for seasonality within years would improve model performance. We used AIC to compare models with and without “month”, selecting the best model for each time series. We extracted the linear slope from the model, which summarises the trend for mean annual temperature change.

To test if biodiversity responses to temperature change were modulated by the baseline climate at any given location, we extracted annual mean temperature data from the WorldClim^53^ database for terrestrial time series, and from the Bio-ORACLE database^54,55^ for marine time series (on a resolution of 0.01° for terrestrial and of 0.1° for marine systems, respectively). For each realm, we standardized the long-term annual mean temperature across all the locations by subtracting the mean and dividing by the standard deviation.

### Meta-analysis

Having estimated the trends for biodiversity and temperature independently for each individual time series, we assessed the effect of temperature change on the rates of change of each biodiversity metric in a second-stage analysis. We employed a meta-analytical Bayesian framework using the package brms^56,57^ (version 2.6.0), and fitted generalized linear models to each realm separately, having initially evaluated that there was an effect of realm when fitting a full model. All models were created using the Stan computational framework (http://mc-stan.org/) accessed via brms. To determine whether the baseline climate modulated the biodiversity responses, models were fit with an interaction term between temperature change and the long-term average temperature at each location. Additionally, we fitted similar models using latitude. We used two random effect terms: one allowing for different slopes per taxonomic group (Taxon), and another allowing for different intercepts per study ID nested within Taxon. This allowed us to account for: 1) potentially different responses to temperature change among taxa; 2) differences in species richness among taxa, as well as different assemblage time series originating from the same study, and different studies monitoring the same taxonomic groups across the BioTIME database, respectively; and 3) spatial autocorrelation. The different taxonomic groups were informed by the original data sources metadata, and were: “Amphibians”, “Benthos”, “Birds”, “Fish”, “Mammals”, “Marine invertebrates”, “Terrestrial invertebrates”, “Terrestrial plants”, and “All - several major groups”.

The overall model structure implemented for each realm was:

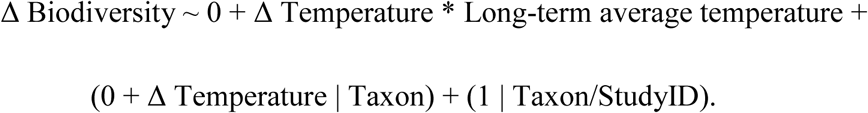

Additionally, brms allows to specify known standard errors for performing meta-analysis using the function se() when specifying the formula for the models^56,57^, using the syntax:

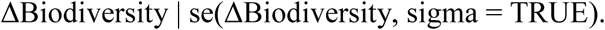

Models were run using 4 chains, each with 8000 iterations, with a warm up of 4000 and non-informative flat priors. Stan implements Hamiltonian Monte Carlo and its extension, the No-U-Turn Sampler (NUTS) algorithms, which converge quickly^56^. Convergence was assessed by visually examining trace plots and using Rhat values (the ratio of the effective sample size to the overall number of iterations, with values close to one indicating convergence)^56^. All the analyses were run in R version 4.3.1^58^.

### Sensitivity analysis

To evaluate the robustness of potential interactions with the baseline climate, we additionally ran our models with two alternative baseline temperature variables. To that end, we extracted the variables “Mean Temperature of Warmest Quarter” from WorldClim and “Long-term maximum sea surface temperature” from Bio-ORACLE, as well as the average temperature in the first year sampled for each biodiversity time series from the same dataset that was used to quantify the trends (i.e. the spatially less resolved HadCRUT4 dataset).

To evaluate whether uneven sampling could be driving the observed differences between the marine and terrestrial realms, we fitted models to subsets of the marine data that matched both the number of locations (1,625 time series) and the latitudinal range of the terrestrial data. We did not attempt to control for temperature change differences between realms because this is part of the signal to be modelled. We fitted the meta-analytical models to 100 random subsamples for each biodiversity metric, illustrating that the estimates for the marine realm are robust (Fig. S4). We further evaluated that biodiversity responses did not show any clear pattern as a function of the number of years sampled, illustrating that the duration of sampling is unlikely to drive our findings (Fig. S6).

We initially explored biodiversity change patterns in response to temperature change using all the assemblage time series with at least five years of sampling – i.e. including tropical and polar locations, yielding 22,119 time series from 179 original studies (Table S1). However, the paucity of tropical and polar locations would prevent us from reliably assessing biodiversity trends in those regions; therefore, we decided to exclude these regions from the analysis and focus on the subtropical and temperate regions, where we have most data.

## Acknowledgements

We are grateful to all the scientists, data collectors and their funders for making data publicly available. We thank the University of St Andrews Bioinformatics Unit (Wellcome Trust ISSF grant 105621/Z/14/Z). L.H.A. acknowledges funding from PBL Netherlands Environmental Assessment Agency, as part of the GLOBIO project (www.globio.info), and from the Jane and Aatos Erkko foundation. C.W. was supported by the Natural Environmental Research Council (grant number 563 NE/L002531/1). M.D. is funded by a Leverhulme Fellowship and by the John Templeton Foundation grant #60501 “Putting the Extended Evolutionary Synthesis to the Test”. The BioTIME database was created using funding from the European Research Council (AdG BioTIME (250198) and ERC PoC BioCHANGE (727440) to A.E.M.).

## Contributions

M.D. and A.M.S. conceived the idea, and all authors contributed to design the project. L.H.A. analysed the data in close consultation with S.A.B., A.E.B., M.D. and A.M.S.. L.H.A. wrote the first draft of the manuscript, with substantial input from A.E.B., M.D. and A.M.S.; all authors contributed to manuscript completion and revision. M.D. and A.M.S. are shared senior authors.

## Data Deposition Statement

All the data can be accessed through the BioTIME database on Zenodo (https://doi.org/10.5281/zenodo.1211105) or through the BioTIME website (http://biotime.standrews.ac.uk/). Code will be made available upon publication.

## References

1. Parmesan, C. & Yohe, G. A globally coherent fingerprint of climate change impacts across natural systems. Nature 421, 37–42 (2003).

2. Parmesan, C. Ecological and Evolutionary Responses to Recent Climate Change. Annu. Rev. Ecol. Evol. Syst. 37, 637–669 (2006).

3. Poloczanska, E. S. et al. Global imprint of climate change on marine life. Nat. Clim. Chang. 3, 919–925 (2013).

4. Bates, A. E. et al. Defining and observing stages of climate-mediated range shifts in marine systems. Glob. Environ. Chang. 26, 27–38 (2014).

5. Chen, I.-C., Hill, J. K., Ohlemüller, R., Roy, D. B. & Thomas, C. D. Rapid range shifts of species of climate warming. Science 333, 1024–1026 (2011).

6. Deutsch, C. A. et al. Impacts of climate warming on terrestrial ectotherms across latitude. Proc. Natl. Acad. Sci. 105, 6668–6672 (2008).

7. Cheung, W. W. L., Watson, R. & Pauly, D. Signature of ocean warming in global fisheries catch. Nature 497, 365–368 (2013).

8. García Molinos, J. et al. Climate velocity and the future global redistribution of marine biodiversity. Nat. Clim. Chang. 6, 83–88 (2016).

9. Sunday, J. M., Bates, A. E. & Dulvy, N. K. Global analysis of thermal tolerance and latitude in ectotherms. Proc. R. Soc. B Biol. Sci. 278, 1823–1830 (2011).

10. Dillon, M. E., Wang, G. & Huey, R. B. Global metabolic impacts of recent climate warming. Nature 467, 704–706 (2010).

11. Comte, L. & Olden, J. D. Climatic vulnerability of the world’s freshwater and marine fishes. Nat. Clim. Chang. 7, 718–722 (2017).

12. Burrows, M. T. et al. The pace of shifting climate in marine and terrestrial ecosystems. Science 334, 652–655 (2011).

13. Darwin, C. R. On the Origin of Species by Means of Natural Selection. (John Murray, 1859).

14. Lenoir, J. et al. Species better track the shifting isotherms in the oceans than on lands. bioRxiv 765776 (2019). doi: 10.1101/765776

15. Pinsky, M. L., Eikeset, A. M., McCauley, D. J., Payne, J. L. & Sunday, J. M. Greater vulnerability to warming of marine versus terrestrial ectotherms. Nature 569, 108–111 (2019).

16. Sunday, J. M., Bates, A. E. & Dulvy, N. K. Thermal tolerance and the global redistribution of animals. Nat. Clim. Chang. 2, 686–690 (2012).

17. Sunday, J. M. et al. Species traits and climate velocity explain geographic range shifts in an ocean-warming hotspot. Ecol. Lett. 18, 944–953 (2015).

18. Burrows, M. T. et al. Ocean community warming responses explained by thermal affinities and temperature gradients. Nat. Clim. Chang. (2019,. in press).

19. Pinsky, M. L., Worm, B., Fogarty, M. J., Sarmiento, J. L. & Levin, S. A. Marine Taxa Track Local Climate Velocities. Science 341, 1239–1242 (2013).

20. Suggitt, A. J. et al. Extinction risk from climate change is reduced by microclimatic buffering. Nat. Clim. Chang. 8, 713–717 (2018).

21. Supp, S. & Ernest, S. Species-level and community-level responses to disturbance: a cross-community analysis. Ecology 95, 1717–1723 (2014).

22. Schipper, A. M. et al. Contrasting changes in the abundance and diversity of North American bird assemblages from 1971) to 2010. Glob. Chang. Biol. 22, 3948–3959 (2016).

23. Dornelas, M. et al. BioTIME: A database of biodiversity time series for the anthropocene. Glob. Ecol. Biogeogr. 27, 760–786 (2018).

24. Blowes, S. A. et al. The geography of biodiversity change in marine and terrestrial assemblages. Science 366, 339–345 (2019).

25. Jones, P. D. et al. Hemispheric and large-scale land-surface air temperature variations: An extensive revision and an update to 2010. J. Geophys. Res. Atmos. 117, D05127 (2012).

26. Harris, I., Jones, P. D., Osborn, T. J. & Lister, D. H. Updated high-resolution grids of monthly climatic observations – the CRU TS3.10 Dataset. Int. J. Climatol. 34, 623–642 (2014).

27. Bowler, D. E. et al. Mapping human pressures across the planet uncovers anthropogenic threat complexes. bioRxiv 432880 (2019). doi: 10.1101/432880

28. Brown, J., Gillooly, J., Allen, A. & Savage, V. Toward a metabolic theory of ecology. Ecology 85, 1771–1789 (2004).

29. Waldock, C., Dornelas, M. & Bates, A. E. Temperature-Driven Biodiversity Change: Disentangling Space and Time. Bioscience 68, 873–884 (2018).

30. Edgar, G. J. et al. Abundance and local-scale processes contribute to multi-phyla gradients in global marine diversity. Sci. Adv. 3, e1700419 (2017).

31. Beaugrand, G., Edwards, M., Raybaud, V., Goberville, E. & Kirby, R. R. Future vulnerability of marine biodiversity compared with contemporary and past changes. Nat. Clim. Chang. 5, 695–701 (2015).

32. Pecl, G. T. et al. Biodiversity redistribution under climate change: Impacts on ecosystems and human well-being. Science 355, eaai9214 (2017).

33. IPCC. Climate Change 2014: Impacts, Adaptation, and Vulnerability. Part A: Global and Sectoral Aspects. Contribution of Working Group II to the Fifth Assessment Report of the Intergovernmental Panel on Climate Change. (Cambridge University Press, 2014).

34. Bates, A. E. et al. Resilience and signatures of tropicalization in protected reef fish communities. Nat. Clim. Chang. 4, 62–67 (2014).

35. Bates, A. E., Stuart-smith, R. D., Barrett, N. S. & Edgar, G. J. Biological interactions both facilitate and resist climate-related functional change in temperate reef communities. Proc. R. Soc. B Biol. Sci. 284, 20170484 (2017).

36. Devictor, V. et al. Differences in the climatic debts of birds and butterflies at a continental scale. Nat. Clim. Chang. 2, 121–124 (2012).

37. Bowler, D. E. et al. Cross-realm assessment of climate change impacts on species’ abundance trends. Nat. Ecol. Evol. 1, 67 (2017).

38. Devictor, V., Julliard, R., Couvet, D. & Jiguet, F. Birds are tracking climate warming, but not fast enough. Proc. R. Soc. B Biol. Sci. 275, 2743–2748 (2008).

39. Menéndez, R. et al. Species richness changes lag behind climate change. Proc. R. Soc. B Biol. Sci. 273, 1465–1470 (2006).

40. Bertrand, R. et al. Ecological constraints increase the climatic debt in forests. Nat. Commun. 7, 12643 (2016).

41. Dornelas, M. et al. Assemblage time series reveal biodiversity change but not systematic loss. Science 344, 296–299 (2014).

42. Meyer, C., Kreft, H., Guralnick, R. & Jetz, W. Global priorities for an effective information basis of biodiversity distributions. Nat. Commun. 6, 8221 (2015).

43. Williams, J. W., Jackson, S. T. & Kutzbach, J. E. Projected distributions of novel and disappearing climates by 2100 AD. Proc. Natl. Acad. Sci. 104, 5738–5742 (2007).

44. Nolan, C. et al. Past and future global transformation of terrestrial ecosystems under climate change. Science 361, 920–923 (2018).

45. Ordonez, A., Williams, J. W. & Svenning, J.-C. Mapping climatic mechanisms likely to favour the emergence of novel communities. Nat. Clim. Chang. 6, 1104–1109 (2016).

46. Doak, D. F. & Morris, W. F. Demographic compensation and tipping points in climate-induced range shifts. Nature 467, 959–962 (2010).

47. Bryndum-Buchholz, A. et al. Twenty-first-century climate change impacts on marine animal biomass and ecosystem structure across ocean basins. Glob. Chang. Biol. 25, 459–472 (2019).

48. IPBES. Report of the Plenary of the Intergovernmental Science-Policy Platform on Biodiversity and Ecosystem Services on the work of its seventh session. (2019).

49. Şekercioğlu, Ç. H., Primack, R. B. & Wormworth, J. The effects of climate change on tropical birds. Biol. Conserv. 148, 1–18 (2012).

50. Chao, A. & Jost, L. Coverage-based rarefaction and extrapolation: standardizing samples by completeness rather than size. Ecology 93, 2533–2547 (2012).

51. Gotelli, N. J. & Colwell, R. K. Quantifying biodiversity: procedures and pitfalls in the measurement and comparison of species richness. Ecol. Lett. 4, 379–391 (2001).

52. Wood, S. N. Generalized Additive Models: An Introduction with R. (Chapman and Hall/CRC, 2017).

53. Fick, S. E. & Hijmans, R. J. WorldClim 2: new 1-km spatial resolution climate surfaces for global land areas. Int. J. Climatol. 37, 4302–4315 (2017).

54. Tyberghein, L. et al. Bio-ORACLE: A global environmental dataset for marine species distribution modelling. Glob. Ecol. Biogeogr. 21, 272–281 (2012).

55. Assis, J. et al. Bio-ORACLE v2.0: Extending marine data layers for bioclimatic modelling. Glob. Ecol. Biogeogr. 27, 277–284 (2018).

56. Bürkner, P.-C. brms: An R Package for Bayesian Multilevel Models Using Stan. J. Stat. Softw. 80, 1–28 (2017).

57. Bürkner, P.-C. Advanced Bayesian Multilevel Modeling with the R Package brms. R J. 10, 395–411 (2018).

58. R Core Team. R: A language and environment for statistical computing. R Foundation for Statistical Computing, Vienna, Austria. http://www.R-project.org/. (2017).

