## Supplementary Information for "Temperature-related biodiversity change across temperate marine and terrestrial systems"

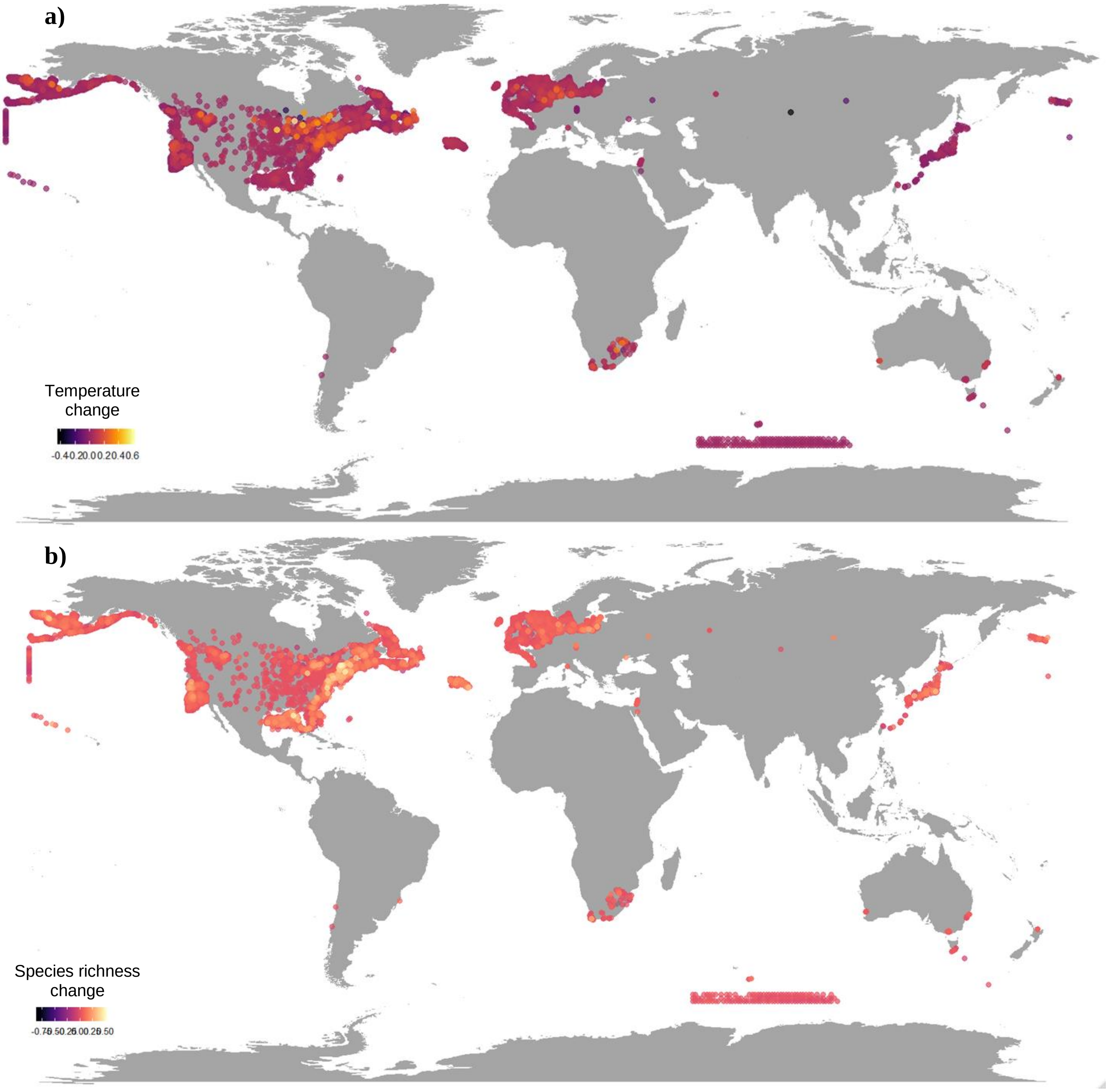


**Figure S1**. Location of temperate biodiversity time series, coloured according to the temperature change experienced during the period of biodiversity monitoring in those locations (a), with the corresponding rates of species richness change (b).

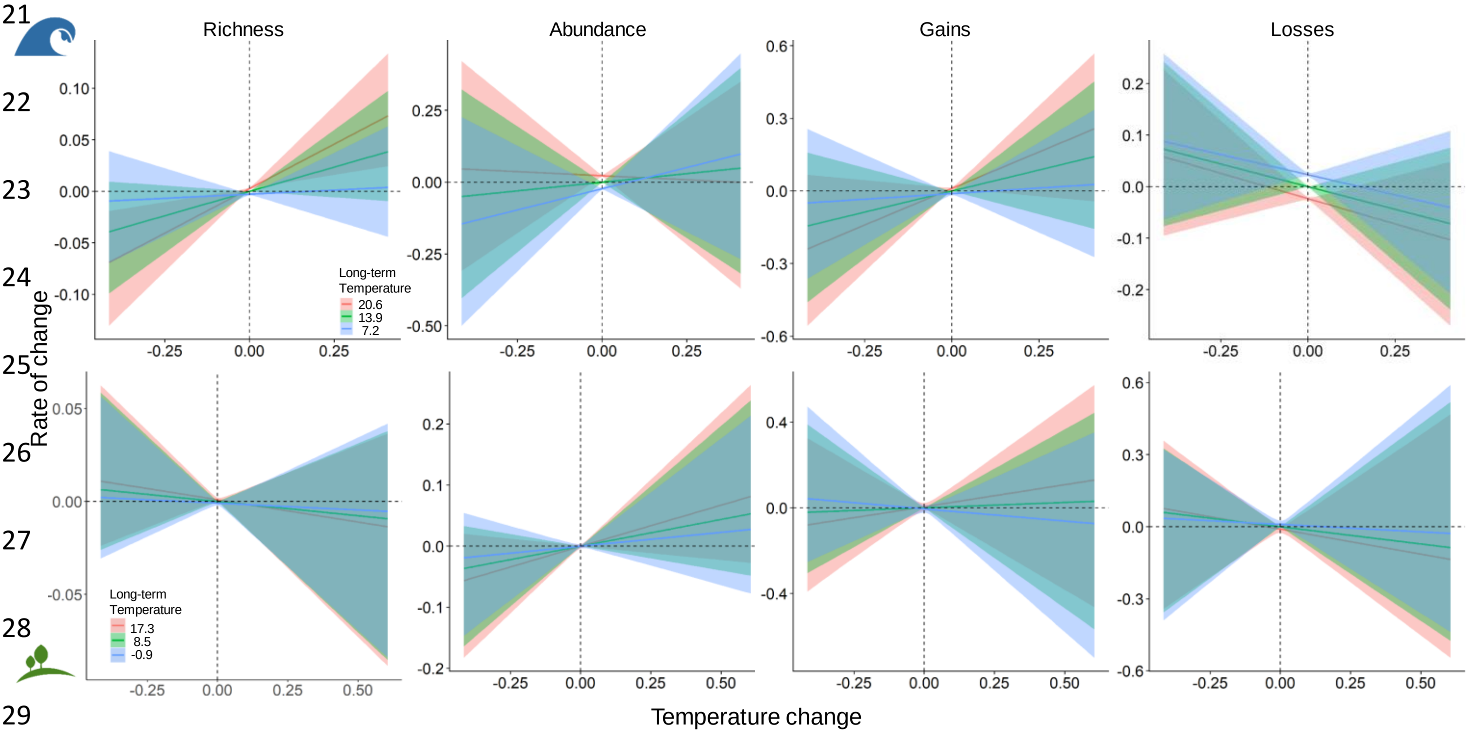


**Figure S2**. Biodiversity responses (rate/year) to the combination of temperature change (°C/year) and long-term mean annual temperature (standardized across all locations) from the meta-analytical models. The coloured fitted lines represent three long-term annual mean temperature values representing the range across the time series (specifically the mean ± one standard deviation for each realm). For each biodiversity metric, the top row is for marine locations, and the bottom row for terrestrial locations (note the different scales among metrics).

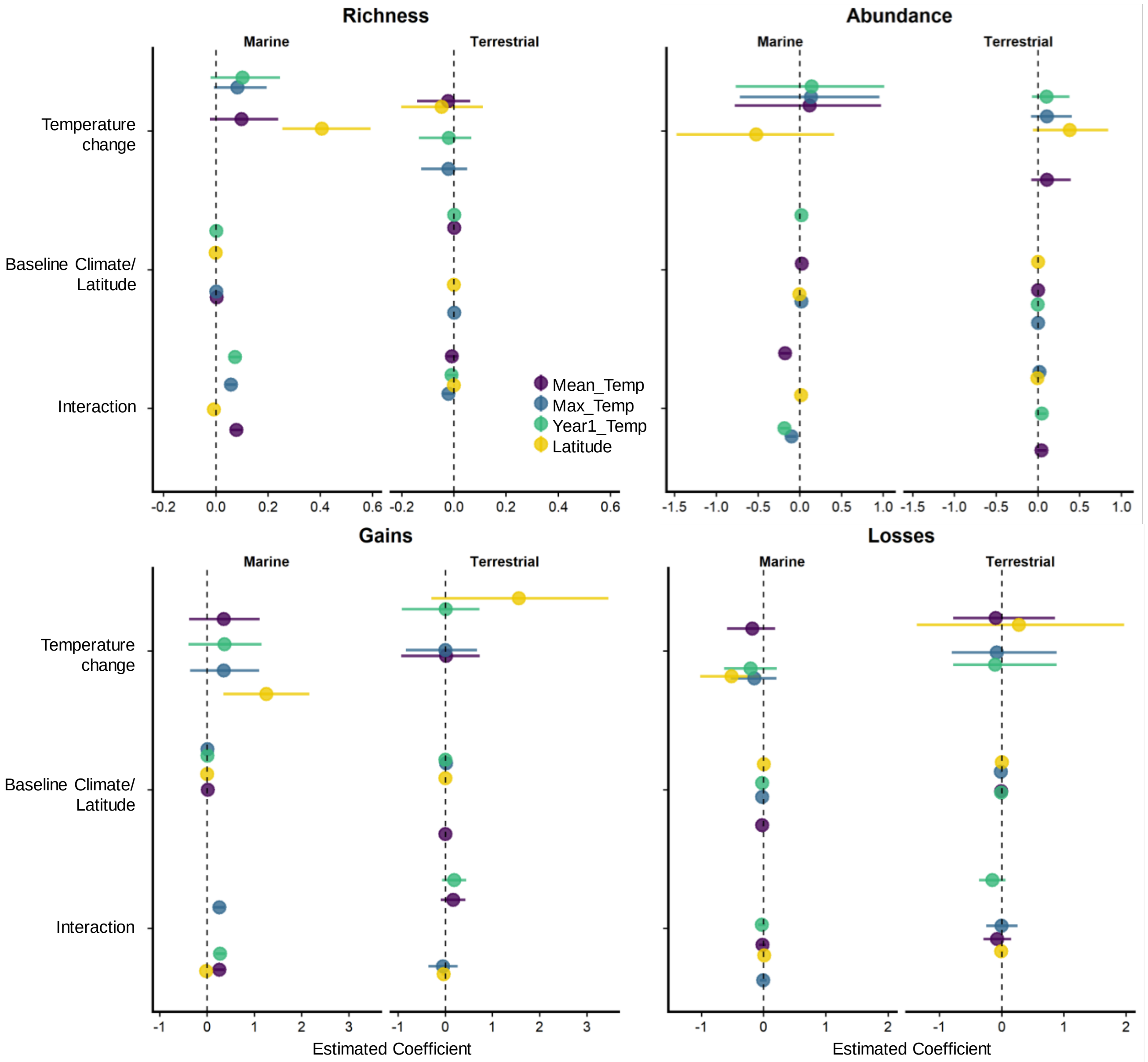


**Figure S3**. Comparison of the meta-analytical model estimates using different variables for baseline climate, as well as latitude, testing for modulating effects of biodiversity responses to temperature change. Our results are overall robust to the different temperature variables used: long-term annual mean and maximum temperature from the databases WorldClim and Bio-ORACLE, and average air and sea surface temperature in the first year sampled from the HadCRUT4 database. Additionally, latitude did not show interacting effects with temperature change.

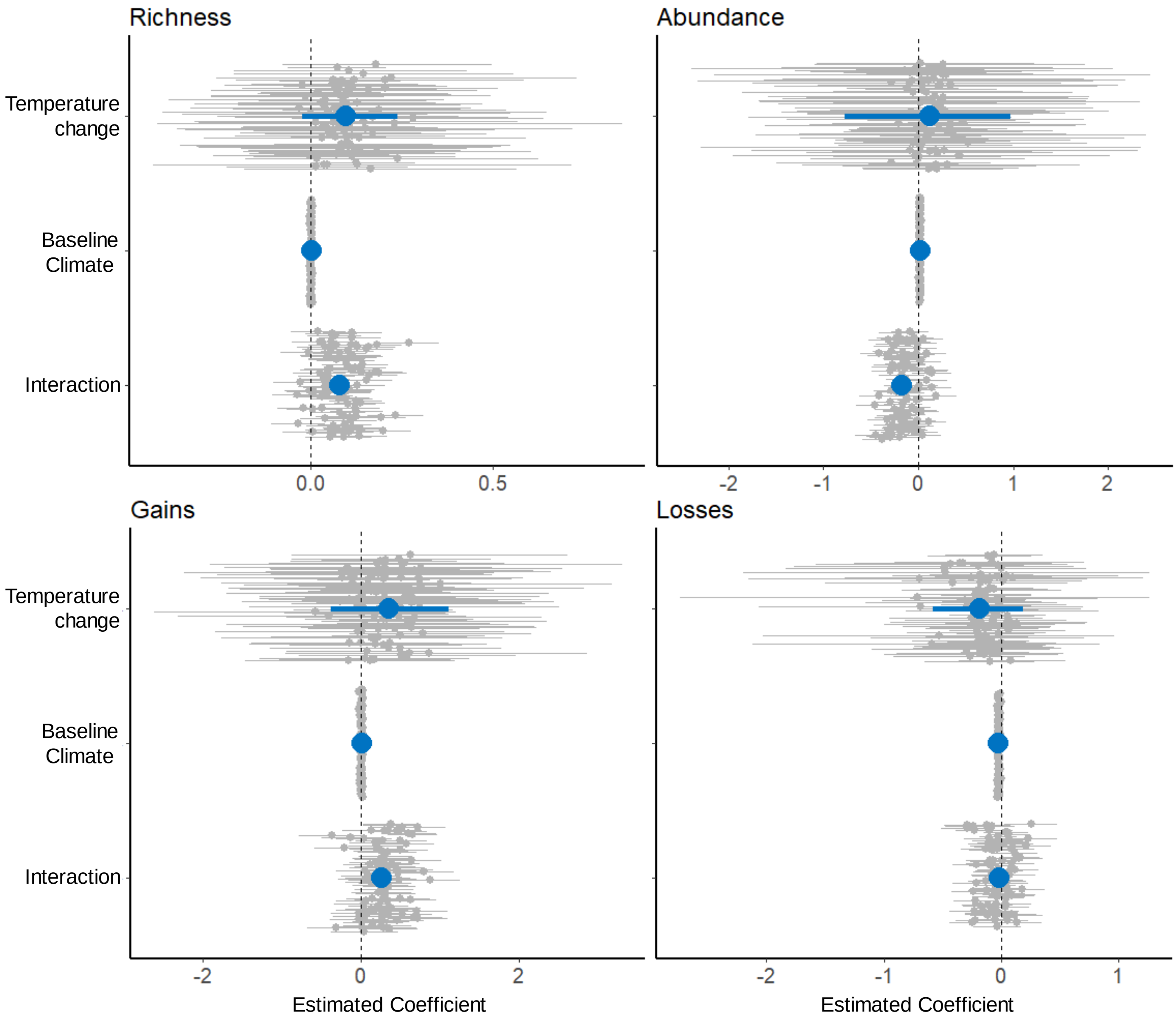


**Figure S4**. Sensitivity analysis for each biodiversity metric. The grey points show the estimated coefficients (and their 95% credible interval) from 100 meta-analytical models fit to subsets of the marine data, which were randomly subsampled to match the number of locations and latitudinal range of the terrestrial data. Despite the increase in uncertainty due to the smaller data subsets (i.e., larger credible intervals), comparing the parameter estimates based on the random sub-samples with the parameters estimated using the entire data (blue dots) shows that the marine estimates were not biased due to uneven sampling.

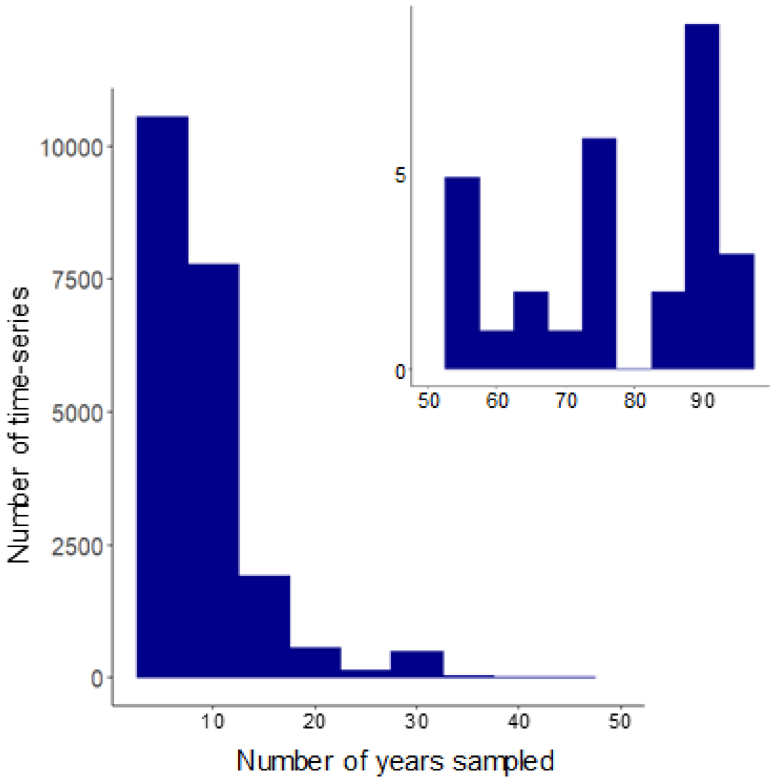


**Figure S5**. Number of years sampled across the time series (inset shows the distribution for time series with >50 years sampled).

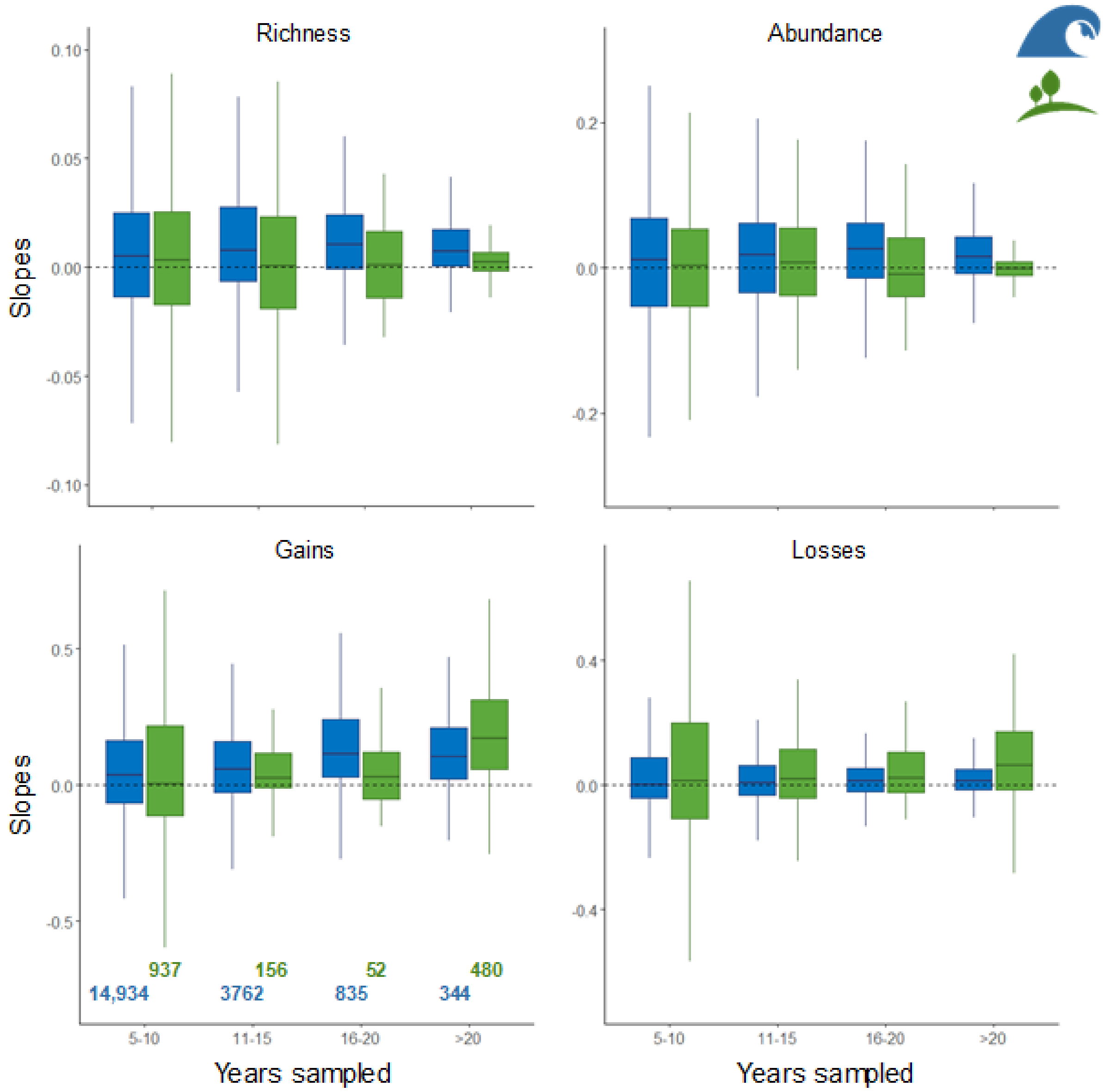


**Figure S6**. Variation in biodiversity estimated slopes as a function of the number of years sampled used to calculate the trends (5-year bins; blue for marine, green for terrestrial); the number of time series in each category is indicated in the panel for gains.

**Table S1**. Biodiversity datasets Study ID and respective citations (* indicates studies not republished with the BioTIME database, but publicly available via the primary references listed).

| STUDY_ID | Ref (s) |
|---|---|
| 18 | 1 |
| 33 | 2, 3 |
| 39 | 4-7 |
| 41 | 8 |
| *42 | 9-15 |
| *44 | 16-18 |
| 45 | 19, 20 |
| 46 | 21 |
| 47 | 22 |
| 51 | 23, 24 |
| 54 | 25 |
| 56 | 26 |
| 58 | 27, 28 |
| 59 | 29 |
| 60 | 30-35 |
| 63 | 36, 37 |
| 67 | 38 |
| 70 | 39 |
| 71 | 40 |
| 78 | 41 |
| 81 | 42, 43 |
| 85 | 44 |
| 86 | 45 |
| 87 | 46-48 |
| 91 | 49 |
| *100 | 50-52 |
| *101 | 50-52 |
| 108 | 53 |
| 112 | 54 |
| 119 | 55 |
| 121 | 56 |
| 123 | 57 |
| 125 | 58 |
| 126 | 59 |
| 127 | 60 |
| 143 | 61, 62 |
| 152 | 63 |
| 163 | 64 |
| 166 | 65-70 |

| | |
|---|---|
| 169 | 71-74 |
| 171 | 75 |
| 172 | 76-79 |
| 176 | 80 |
| 180 | 81 |
| 182 | 82 |
| 183 | 83 |
| 189 | 84 |
| 190 | 85 |
| 191 | 86 |
| 192 | 87 |
| *193 | 50-52 |
| 194 | 88 |
| 195 | 89 |
| 196 | 90 |
| *197 | 91 |
| *198 | 92 |
| 200 | 86 |
| 204 | 93 |
| *205 | 94 |
| *206 | 95 |
| *207 | 96 |
| *208 | 97 |
| *209 | 98 |
| *210 | 99 |
| 211 | 100 |
| 212 | 101 |
| 213 | 102 |
| 214 | 103 |
| *215 | 104 |
| *216 | 105 |
| 217 | 106 |
| *218 | 107 |
| *219 | 108 |
| *220 | 109 |
| 221 | 110 |
| 225 | 111 |
| 231 | 112 |
| 232 | 113 |
| 234 | 114-120 |
| 240 | 121 |
| 242 | 122 |
| 243 | 123, 124 |
| *244 | 125 |
| 246 | 126 |
| 249 | 127 |

| | |
|---|---|
| 252 | 128 |
| 255 | 129 |
| *256 | 98 |
| 271 | 130 |
| 272 | 131 |
| 273 | 132 |
| 275 | 133 |
| *277 | 16-18 |
| *279 | 16-18 |
| 287 | 57 |
| 288 | 55 |
| 294 | 134, 135 |
| 295 | 136 |
| 296 | 136 |
| 297 | 137 |
| 300 | 138 |
| 301 | 139, 140 |
| 305 | 141 |
| 308 | 142 |
| *309 | 143 |
| 310 | 144 |
| 311 | 145 |
| 312 | 146 |
| 313 | 147 |
| 316 | 148 |
| 317 | 149 |
| *318 | 150 |
| 319 | 151 |
| 321 | 152 |
| 325 | 153 |
| 327 | 154 |
| 329 | 155 |
| 336 | 29 |
| 339 | 156 |
| 340 | 157 |
| 348 | 158 |
| 354 | 159 |
| 356 | 160 |
| 357 | 161 |
| 358 | 162 |
| 359 | 163 |
| 361 | 164 |
| 363 | 165 |
| 365 | 166 |
| 366 | 167 |
| 367 | 168 |

| | |
|---|---|
| 369 | 169 |
| 372 | 170 |
| 373 | 171 |
| 374 | 172 |
| 375 | 173 |
| 377 | 174 |
| 378 | 175 |
| 379 | 176 |
| 380 | 177, 178 |
| 381 | 179 |
| 382 | 177, 180 |
| 412 | 181 |
| 413 | 182 |
| 414 | 182 |
| 415 | 182 |
| 416 | 182 |
| 419 | 183 |
| 420 | 184 |
| 422 | 185 |
| 423 | 185 |
| 424 | 186 |
| 428 | 187-190 |
| 435 | 191 |
| 436 | 192 |
| 438 | 193 |
| 439 | 194 |
| 440 | 194 |
| 441 | 194 |
| 442 | 195 |
| 444 | 196 |
| 447 | 197 |
| 461 | 198-200 |
| 462 | 198-200 |
| 463 | 198-200 |
| 464 | 198-200 |
| 465 | 198-200 |
| 466 | 201 |
| 468 | 202 |
| 469 | 203 |
| 471 | 204 |
| 473 | 205 |
| 475 | 206 |
| 477 | 207, 208 |
| 499 | 209 |
| 500 | 210 |
| 502 | 211 |

| | |
|---|---|
| 504 | 212 |
| 505 | 213 |

110

**Table S2**. Coefficients estimated from the Bayesian meta-analytical models for each biodiversity metric in each realm, showing the Posterior Mean Estimates and the lower and upper 95% Credible Intervals.

| Biodiversity Metric | Realm | Variable | Posterior Mean Estimate | Lower 95% Cred. Interval | Upper 95% Cred. Interval | Rhat |
|---|---|---|---|---|---|---|
| Species Richness | Marine | Temperature change | 0.098 | -0.023 | 0.239 | 1.0013 |
| | | Long-term Average Temp | 0.003 | 0.002 | 0.004 | 1.0000 |
| | | Interaction | 0.079 | 0.057 | 0.1004 | 0.9999 |
| | Terrestrial | Temperature change | -0.023 | -0.141 | 0.063 | 1.0018 |
| | | Long-term Average Temp | 0.001 | 0.000 | 0.002 | 0.9998 |
| | | Interaction | -0.007 | -0.028 | 0.013 | 1.0001 |
| Abundance | Marine | Temperature change | 0.117 | -0.778 | 0.973 | 1.0005 |
| | | Long-term Average Temp | 0.023 | 0.019 | 0.026 | 0.9998 |
| | | Interaction | -0.177 | -0.237 | -0.116 | 0.9999 |
| | Terrestrial | Temperature change | 0.107 | -0.080 | 0.394 | 1.0010 |
| | | Long-term Average Temp | 0.000 | -0.003 | 0.003 | 1.0002 |
| | | Interaction | 0.043 | -0.013 | 0.099 | 1.0000 |
| Species Gains | Marine | Temperature change | 0.353 | -0.383 | 1.108 | 1.0011 |
| | | Long-term Average Temp | 0.011 | 0.005 | 0.017 | 1.0000 |
| | | Interaction | 0.256 | 0.148 | 0.362 | 0.9999 |
| | Terrestrial | Temperature change | 0.015 | -0.936 | 0.733 | 1.0003 |
| | | Long-term Average Temp | 0.005 | -0.013 | 0.023 | 1.0000 |
| | | Interaction | 0.167 | -0.095 | 0.430 | 0.9998 |
| Species Losses | Marine | Temperature change | -0.183 | -0.585 | 0.185 | 1.0009 |
| | | Long-term Average Temp | -0.023 | -0.027 | -0.020 | 0.9998 |
| | | Interaction | -0.019 | -0.076 | 0.038 | 0.9999 |
| | Terrestrial | Temperature change | -0.095 | -0.784 | 0.857 | 1.0006 |
| | | Long-term Average Temp | -0.010 | -0.025 | 0.005 | 1.0001 |
| | | Interaction | -0.075 | -0.297 | 0.152 | 1.0005 |

### Data sources references


1. Zachmann, L., Moffet, C. & Adler, P. Mapped quadrats in sagebrush steppe: long-term data for analyzing demographic rates and plant–plant interactions. *Ecology*, **91**, 3427 (2010).
2. Widdicombe, C. E., Eloire, D., Harbour, D., Harris, R. P. & Somerfield, P. J. Long-term phytoplankton community dynamics in the Western English Channel. *Journal of Plankton Research*, **32**, 643–655 (2010). doi:10.1093/plankt/fbp127
3. Widdicombe, C. E., Eloire, D., Harbour, D., Harris, R. P. & Somerfield, P. J. Time series of phyto- and microzooplankton abundance and composition at station L4 in the English Channel from 1988 to 2009 (2010). doi:10.1594/PANGAEA.758061
4. Holmes, R. T. & Sherry, T. W. Assessing population trends of New Hampshire forest birds: Local versus regional patterns. *The Auk*, **105**, 756–768 (1988).
5. Holmes, R. T. & Sherry, T. W. Thirty-year bird population trends in an unfragmented temperate deciduous forest: the importance of habitat change. *The Auk*, **118**, 589–610 (2001).
6. Holmes, R. T. & Sturges, F. W. Bird community dynamics and energetics in a northern hardwoods ecosystem. *Journal of Animal Ecology*, **44**, 175–200 (1975).
7. Holmes, R. T., Sherry, T. W. & Sturges, F. W. Bird Community Dynamics in a Temperate Deciduous Forest: Long-Term Trends at Hubbard Brook. *Ecological Monographs*, **56**, 201–220 (1986).
8. Preston, F. W. Time and Space and the Variation of Species. *Ecology*, **41**, 611–627 (1960).
9. B. Stone *et al*., Population estimates of birds in Britain and in the United Kingdom. *British Birds*, **90**, 1 (1997).

10. D. W. Gibbons, J. B. Reid, R. A. Chapman, The New Atlas of Breeding Birds in Britain and Ireland: 1988-1991 (1994).
11. G. Beven, Changes in breeding bird populations of an oak-wood on Bookham Common, Surrey, over twenty-seven years. *London Naturalist,* **55**, 23 (1976).
12. K. J. Gaston, T. M. Blackburn, Pattern and process in macroecology. Wiley-Blackwell, Oxford, England (2000).
13. M. Williamson, Are communities ever stable? Colonization, succession and stability. AJ Gray, MJ Crawley & PJ Edwards (Eds.), Blackwell Scientific Publications, Oxford, (1987) pp 353.
14. P. Lack, The atlas of wintering birds in Britain and Ireland. A&C Black, London, England (2010).
15. P. Standley, A. Swash, R. Gillmor, The birds of Berkshire. Berkshire Atlas Group, Berkshire, England (1996).
16. Halpern, C. B. & Dyrness, C. "Plant succession and biomass dynamics following logging and burning in the Andrews Experimental Forest Watersheds 1 and 3, 1962-Present". Long-Term Ecological Research. Forest Science Data Bank, Corvallis. **Available at: http://andrewsforest.oregonstate.edu/data/abstract.cfm?dbcode=TP073**, accessed 2012 (2010).
17. Halpern, C. B. & Lutz, J. A. "Canopy closure exerts weak controls on understory dynamics: a 30-year study of overstory–understory interactions." **Available at: Dryad DigitalRepository doi:10.5061/dryad.1q88j**, accessed 2013 (2013).
18. Halpern, C.B. & Lutz, J.A. Canopy closure exerts weak controls on understory dynamics: a 30-year study of overstory–understory interactions. *Ecological Monographs*, **83**, 221–237 (2013).

19. Brooks, A.J. "MCR LTER: Coral Reef: Long-term Population and Community Dynamics: Fishes". Moorea Coral Reef. **Available at: http://mcr.lternet.edu/cgi-bin/showDataset.cgi?docid=knb-lter-mcr.6**, accessed 2012.
20. Brooks, A.J. Moorea Coral Reef LTER: Coral Reef: Long-term Population and Community Dynamics: Fishes. **Available at: knb-lter-mcr.6.54 doi:10.6073/pasta/d688610e536f54885a3c59d287f6c4c3**, accessed 2016 (2016).
21. Williamson, M. The Land-Bird Community of Skokholm: Ordination and Turnover. *Oikos*, **41**, 378–384 (1983).
22. Vickery, W. L. & Nudds, T. D. Detection of Density-Dependent Effects in Annual Duck Censuses. *Ecology*, **65**, 96 (1984).
23. "Fluctuations and long-term trends in the relative densities of tetraonid populations in Finland, 1964-77." NERC Centre for Population Biology, Imperial College. The Global Population Dynamics Database v2.0. **Available at: https://www.imperial.ac.uk/cpb/gpdd2/secure/register.aspx**, accessed 2012.
24. Lindén, H. & Rajala, P. Fluctuations and long-term trends in the relative densities of tetraonid populations in Finland, 1964-77. *Finnish Game Research*, **39**, 13-34 (1981).
25. Willig, M. R. & Bloch, C. P. "El Verde Grid long-term invertebrate data: Luquillo Long Term Ecological Research Site Database: Data Set 107". **Available at:http://luq.lternet.edu/data/luqmetadata107/7427**, accessed 2016 (2016).
26. Friggens, M. "Sevilleta LTER Small Mammal Population Data", Albuquerque, NM: Sevilleta Long Term Ecological Research Site Database: SEV008**. Available at: http://sev.lternet.edu/data/sev-8**, accessed 2012 (2008).
27. Waide, R. B. "Bird abundance - point counts, El Verde Field Station, Puerto Rico: Luquillo Long Term Ecological Research Site Database: Data Set 23". **Available at: http:/luq.lternet.edu/data/luqmetadata23**, accessed 2012.

28. Waide, R. B. “Bird abundance - point counts”. Long Term Ecological Research Network. **Available at: http://dx.doi.org/10.6073/pasta/0d96957379936a038ebbbcc6135b2fab**, accessed 2012 (2010).

29. Ernest, S., Valone, T.J. & Brown, J.H. Long-term monitoring and experimental manipulation of a Chihuahuan Desert ecosystem near Portal, Arizona, USA. *Ecology*, **90**, 1708-1708 (2009).

30. Condit, R. Tropical forest census plots: Methods and results from Barro Colorado Island, Panama and a Comparison with other plot. Springer Verlag and RG Landes Company, Berlin (1998).

31. Condit, R., Ashton, P., Bunyavejchewin, S., Dattaraja, H., Davies, S., Esufali, S., Ewango, C., Foster, R., Gunatilleke, I. & Gunatilleke, C. The importance of demographic niches to tree diversity. *Science*, **313**, 98-101 (2006).

32. Condit, R., Chisholm, R.A. & Hubbell, S.P. Thirty years of forest census at Barro Colorado and the importance of immigration in maintaining diversity. *PloS one*, **7**, e49826 (2012).

33. Condit, R., Lao, S., Pérez, R., Dolins, S.B., Foster, R. & Hubbell, S. Dataset: Barro Colorado Forest Census Plot Data (Version 2012) (2012).

34. Condit, R., Pérez, R., Aguilar, S., Lao, S., Robin, F. & Hubbell, S. Tree species abundance through time in tropical forest census plots, Panama. DataONE Dash, Dataset, **Available at: https://doi.org/10.15146/R3MM4V** (2018).

35. Hubbell, S. P., Condit, R. & Foster, R. B. "Barro Colorado Forest Census Plot Data". **Available at: https://ctfs.arnarb.harvard.edu/webatlas/datasets/bci**, accessed 2012 (2005).

36. Moore, N. The development of dragonfly communities and the consequences of territorial behaviour: A 27 year study on small ponds at Woodwalton Fen, Cambridgeshire, United Kingdom. *Odonatologica*, **20**, 203–231 (1991).

37. Moore, N. W. "The development of dragonfly communities and the consequences of territorial behaviour: A 27-year study on small ponds at Woodwalton Fen, Cambridgeshire, United Kingdom". NERC Centre for Population Biology, Imperial College. The Global Population Dynamics Database Version 2.0. **Available at: http://www3.imperial.ac.uk/cpb/databases/gpdd**, accessed 2012 (1991).

38. "Animal Demography Unit - Coordinated Waterbird Counts (CWAC) - AfrOBIS". **Available at http://www.iobis.org/**, accessed 2012.

39. Vanholder, B. "Belgian Migrating Lepidoptera". NERC Centre for Population Biology, Imperial College. The Global Population Dynamics Database v2.0. **Available at: https://www.imperial.ac.uk/cpb/gpdd2/secure/register.aspx**, accessed 2012 (1997).

40. Ratkova, T. N. "Phytoplankton of the White Sea, Barents Sea, Amundsen & Nansen Basins". Institute of Oceanology, Academy of Sciences of Russia, Moscow, Russia; Arctic Ocean Diversity, University of Alaska Fairbanks, Fairbanks. **Available at: http://www.arcodiv.org/Database/Plankton_datasets**, accessed 2012.

41. Zettler, M.L. Macrozoobenthos baltic sea (1980-2005) as part of the IOW-Monitoring. Institut für Ostseeforschung Warnemünde, Germany. **Available at: http://www.iobis.org/mapper/?dataset=2289**, accessed 2012 (2005).

42. Robinson, K. P. "CRRU (Cetacean Research and Rescue Unit) Cetacean sightings in Scotland waters". **Available at: http://www.emodnet-biology.eu/component/imis/?module=dataset&dasid=2819**, accessed 2012 (2010).

43. Robinson, K.P., Baumgartner, N., Eisfeld, S.M., Clark, N.M., Culloch, R.M., Haskins, G.N., Zapponi, L., Whaley, A.R., Weare, J.S. & Tetley, M.J. The summer distribution and occurrence of cetaceans in the coastal waters of the outer southern Moray Firth in northeast Scotland (UK). *Lutra*, **50**, 19 (2007).

44. Addinck, W. & de Kluijver, M. North Sea observations of Crustacea, Polychaeta, Echinodermata, Mollusca and some other groups between 1986 and 2003. Expert Centre for Taxonomic Idenditification (ETI), the Netherlands. **Available at: http://www.emodnet-biology.eu/data-catalog?module=dataset&dasid=1037**, accessed 2012 (2003).

45. Derezuyk, N. "Phytoplankton of the Ukrainian Black Sea shelf (1985-2005)". **Available at: http://www.emodnet-biology.eu/component/imis/?module=dataset&dasid=2694**, accessed 2012.

46. Bakker, C. & Herman, P.M.J. Phytoplankton in the Oosterschelde before, during and after the storm-surge barrier (1982-1990). Netherlands Institute of Ecology; Centre for Estuarine and Marine Ecology, Netherlands. EurOBIS Data. **Available at: http://www.iobis.org/mapper/?dataset=505**, accessed 2013 (1990).

47. Bakker, C., Herman, P. & Vink, M. A new trend in the development of the phytoplankton in the Oosterschelde (SW Netherlands) during and after the construction of a storm-surge barrier. The Oosterschelde Estuary (The Netherlands): a Case-Study of a Changing Ecosystem, pp. 79-100. Springer (1994).

48. Bakker, C., Herman, P.M.J. & Vink, M. Changes in seasonal succession of phytoplankton induced by the storm-surge barrier in the Oosterschelde (S.W. Netherlands). *Journal of Plankton Research*, **12**, 947-972 (1990).

49. "Baltic Seabirds Transect Surveys", Institute of Ecology of Vilnius University - OBIS-SEAMAP. **Available at: http://www.emodnet-biology.eu/component/imis/?module=dataset&dasid=1971**, accessed 2012.
50. P. A. Henderson, A. E. Magurran, Direct evidence that density-dependent regulation underpins the temporal stability of abundant species in a diverse animal community. *Proceedings of the Royal Society B: Biological Sciences*, **281**, 20141336 (2014).
51. P. A. Henderson, A. E. Magurran. Data from: Direct evidence that density-dependent regulation underpins the temporal stability of abundant species in a diverse animal community. **Available at: http://dx.doi.org/10.5061/dryad.3090c**, accessed 2014 (2014).
52. P. Henderson, The long-term study of the fish and crustacean community of the Bristol Channel. **Available at http://www.pisces-conservation.com/,** accessed 2013.
53. Woehler, E. "Seabirds of the Southern and South Indian Ocean - Australian Antarctic Data Centre". **Available at: http://www.iobis.org**, accessed 2012.
54. "South Western Pacific Regional OBIS Data Asteroid Subset", NIWA (National Institute of Water and Atmospheric Research - New Zealand) MBIS (Marine Biodata Information System) accessed through South Western Pacific OBIS. **Available at: http://www.iobis.org/mapper/?dataset=219**, accessed 2012.
55. Clark, D. & Branton, B. DFO Maritimes Research Vessel Trawl Surveys, OBIS Canada Digital Collections. Bedford Institute of Oceanography, Dartmouth, Nova Scotia, Canada, OBIS Canada (2007).
56. "CRED Towed-Diver Fish Biomass Surveys in the Pacific Ocean 2000-2010". Coral Reef Ecosystem Division (CRED), Pacific Island Fisheries Sciences Center, National Marine Fisheries Service. **Available at: http://www.iobis.org/mapper/?dataset=1581**, accessed 2012 (2011).

57. Sherman, S. "Maine Department of Marine Resources Inshore Trawl Survey, 2000 – 2009". Maine Department of Marine Resources, Maine. **Available at: http://www.usgs.gov/obis-usa/data_search_and_access/datasets.html**, accessed 2012 (2010).

58. Reichert, M. "MARMAP Chevron Trap Survey 1990-2009". SCDNR/NOAA MARMAP Program, SCDNR MARMAP Aggregate Data Surveys, The Marine Resources Monitoring, Assessment, and Prediction (MARMAP) Program, Marine Resources Research Institute, South Carolina Department of Natural Resources U.S.A.. **Available at: http://www.usgs.gov/obis-usa/data_search_and_access/participants.html**, accessed 2012 (2009).

59. Reichert, M. "MARMAP Neuston Nets 1990-2009". SCDNR/NOAA MARMAP Program, SCDNR MARMAP Aggregate data surveys, The Marine Resources Monitoring, Assessment, and Prediction (MARMAP) Program, Marine Resources Research Institute, South Carolina Department of Natural Resources U.S.A.. **Available at: http://www.usgs.gov/obis-usa/data_search_and_access/participants.html**, accessed 2012 (2010).

60. Reichert, M. "MARMAP Florida Antillean Trap Survey 1990-2009". SCDNR/NOAA MARMAP Program, SCDNR MARMAP Aggregate Data Surveys, The Marine Resources Monitoring, Assessment, and Prediction (MARMAP) Program, Marine Resources Research Institute, South Carolina Department of Natural Resources U.S.A.. **Available at: http://www.usgs.gov/obis-usa/data_search_and_access/participants.html**, accessed 2012 (2009).

61. Escribano, R., Manríquez, K. & Godoy, F. "Copepoda-COPAS Center (COPAS_CPD1) - Planktonic copepods from the Chilean Humboldt Current System -

Eastern South Pacific Regional Node of OBIS (ESPOBIS)". **Available at: http://www.iobis.org**, accessed 2012 (2006).

62. Hidalgo, P., Escribano, R., Vergara, O., Jorquera, E., Donoso, K. & Mendoza, P. Patterns of copepod diversity in the Chilean coastal upwelling system. *Deep Sea Research Part II: Topical Studies in Oceanography*, **57**, 2089–2097 (2010).
63. "CMarZ (Census of Marine Zooplankton)-Asia Database". Accessed through OBIS-SCAR-MarBIN. **Available at: http://www.iobis.org/mapper/?dataset=1500**, accessed 2012.
64. "The Observer Program database", accessed through the OBIS-USA North Pacific Groundfish Observer (North Pacific Research Board). **Available at: http://www.iobis.org**, accessed 2012.
65. "PIROP Northwest Atlantic 1965-1992 - OBIS SEAMAP". **Available at: http://www.iobis.org/mapper/?dataset=2245**, accessed 2012.
66. Brown, R. G., Nettleship, D. N., Germain, P., Tull, C. E. & Davis, T. Atlas of eastern Canadian seabirds (1975).
67. Diamond, A., Gaston, A. & Brown, R. Converting PIROP Counts of Seabirds at Sea to Absolute Densities. Progress Notes No 164. Canadian Wildlife Service, Ottawa (1986).
68. Halpin, P. N., Read, A .J., Fujioka, E., Best, B. D., Donnelly, B., Hazen, L. J., Kot, C., Urian, K., LaBrecque, E. & Dimatteo, A. OBIS-SEAMAP: The world data center for marine mammal, sea bird, and sea turtle distributions. *Oceanography*, **22**, 104-115 (2009).
69. Huettmann, F. An ecological GIS research application for the northern Atlantic-The PIROP database software, environmental data sets and the role of the internet. In: Riekert W.-F. and Tochtermann K. (Eds.) Hypermedia im Umweltschutz Proceedings

of Deutsche Gesellschaft für Informatik (GI) and Forschungsinstitut für anwendungsorientierte Wissensverarbeitung (FAW) Ulm. Umwelt-Informatik aktuell; Bd.17, Metropolis Verlag/Marburg. pp. 213-217 (1998).

70. Read, A., Halpin, P., Crowder, L., Best, B. & Fujioka, E. OBIS-SEAMAP: mapping marine mammals, birds and turtles. World Wide Web electronic publication. http://seamap. env. duke. Edu (2010).

71. Jahncke, J. & Rintoul, C. "CalCOFI and NMFS Seabird and Marine Mammal Observation Data, 1987-2006". California Cooperative Oceanic Fisheries Investigations (CalCOFI) and National Marine Fisheries Service (NMFS) cruises, 1987-2006 - OBIS SEAMAP. **Available at: http://www.iobis.org**, accessed 2012 (2006).

72. Rintoul, C., Schlagenhauf-Langabeer, B., Hyrenbach, K. D., Morgan, K. H. & Sydeman, W. J. Atlas of California Current Marine Birds and Mammals: Version 1. Unpublished report, PRBO Conservation Science, Petaluma, California (2006).

73. Yen, P. P. W., Sydeman, W. J. & Hyrenbach, K. D. Marine bird and cetacean associations with bathymetric habitats and shallow-water topographies: Implications for trophic transfer and conservation. *Journal of Marine Systems,* **50**, 79–99 (2004).

74. Yen, P. P. W., Sydeman, W. J., Bograd, S. J. & Hyrenbach, K. D. Spring-time distributions of migratory marine birds in the southern California Current: Oceanic eddy associations and coastal habitat hotspots over 17 years. *Deep-Sea Research Part II: Topical Studies in Oceanography,* **53**, 399–418 (2006).

75. "Bahamas Marine Mammal Research Organisation Opportunistic Sightings - OBIS SEAMAP". **Available at: http://www.iobis.org**, accessed 2012.

76. "POPA cetacean, seabird, and sea turtle sightings in the Azores area 1998-2009 - OBIS SEAMAP". **Available at: http://www.iobis.org/mapper/?dataset=4257**, accessed 2012.

77. Amorim, P., Figueiredo, M., Machete, M., Morato, T., Martins, A. & Serrão Santos, R. Spatial variability of seabird distribution associated with environmental factors: a case study of marine Important Bird Areas in the Azores. *ICES Journal of Marine Science*, **66**, 29-40 (2008).

78. Machete, M. & Santos, R. Azores Fisheries Observer Program (POPA): a case study of the multidisciplinary use of observer data. Proceedings of the 5th International Fisheries Observer Conference, pp. 15-18 (2007).

79. Morato, T., Varkey, D. A., Damaso, C., Machete, M., Santos, M., Prieto, R., Santos, R. S. & Pitcher, T. J. Evidence of a seamount effect on aggregating visitors. *Marine Ecology Progress Series*, **357**, 23-32 (2008).

80. Kennedy, M. & Spry, J. Atlantic Zone Monitoring Program Maritimes Region plankton datasets. Fisheries and Oceans Canada-BioChem archive. OBIS Canada, Bedford Institute of Oceanography, Dartmouth, Nova Scotia, Canada (2011).

81. "East Coast North America Strategic Assessment Project, Groundfish Atlas for the East Coast of North America". **Available at: http://www.iobis.org**, accessed 2012.

82. Wade, E. Snow crab research trawl survey database (Southern Gulf of St. Lawrence, Gulf region, Canada) from 1988 to 2010. OBIS Canada, Bedford Institute of Oceanography, Dartmouth, Nova Scotia, Canada (2011).

83. Tremblay, J. M. & Branton, B. DFO Maritimes Research Vessel Trawl Surveys, OBIS Canada Digital Collections. Bedford Institute of Oceanography, Dartmouth, Nova Scotia, Canada, OBIS Canada (2007).

84. "St. John, USVI Fish Assessment and Monitoring Data (2002 - Present)". Silver Spring, MD Publisher: NOAAs Ocean Service, National Centers for Coastal Ocean Science (NCCOS). National Oceanic and Atmospheric Association (NOAA)-National Ocean Service (NOS)-National Centers for Coastal Ocean Science (NCCOS)-Center for Coastal Monitoring and Assessment (CCMA)-Biogeography Team. **Available at: http://www.iobis.org/mapper/?dataset=1672**, accessed 2012 (2007).

85. "St. Croix, USVI Fish Assessment and Monitoring Data (2002 - Present)". Silver Spring, MD Publisher: NOAAs Ocean Service, National Centers for Coastal Ocean Science (NCCOS). National Oceanic and Atmospheric Association (NOAA)-National Ocean Service (NOS)-National Centers for Coastal Ocean Science (NCCOS)-Center for Coastal Monitoring and Assessment (CCMA)-Biogeography Team**. Available at: http://www.iobis.org/mapper/?dataset=1673**, accessed 2012 (2007).

86. "NEFSC Benthic Database (OBIS-USA)". Northeast Fisheries Science Center, National Marine Fisheries Service, NOAA, U.S. Department of Commerce. **Available at: http://www.iobis.org/mapper/?dataset=1694**, accessed 2012 (2010).

87. "Whale Catches in Southern Ocean". OBIS - Australian Antarctic Data Centre. **Available at: http://www.iobis.org**, accessed 2013.

88. Jones, J. & Miller, J. "Spatial and temporal distribution and abundance of moths in the Andrews Experimental Forest, 1994 to 2008". H. J. Andrews Experimental Forest. Forest Science Data Bank, Corvallis. **Available at: http://andrewsforest.oregonstate.edu/data/abstract.cfm?dbcode=SA015**, accessed 2013.

89. USGS Patuxent Wildlife Research Center “North American Breeding Bird Survey” ftp data set, version 2014.0. **Available at: ftp://ftpext.usgs.gov/pub/er/md/laurel/BBS/DataFiles/**, accessed 2013.

90. Moore, J. J. & Howson, C. M. “Survey of the rocky shores in the region of Sullom Voe, Shetland, A report to SOTEAG from Aquatic Survey & Monitoring Ltd”, Cosheston, Pembrokeshire. 29 p. **Available at: http://www.soteag.org.uk**, accessed 2013.
91. DATRAS "Scottish West Coast Survey For Commercial Fish Species 1985-2013". **Available at https://datras.ices.dk**, accessed 2013.
92. DATRAS "ICES Baltic International Trawl Survey For Commercial Fish Species (1991 - 2013)". **Available at https://datras.ices.dk**, accessed 2013.
93. Degraer, S., Wittoeck, J., Appeltans, W., Cooreman, K., Deprez, T., Hillewaert, H., Hostens, K., Mees, J., Vanden Berghe, E. & Vincx, M. “Macrobel: Long term trends in the macrobenthos of the Belgian Continental Shelf.” Oostende, Belgium. **Available at: http://www.emodnet-biology.eu/data-catalog?module=dataset&dasid=145**, accessed 2013 (2006).
94. DATRAS “Fish trawl survey: Scottish Rockall Survey for commercial fish species. ICES Database of trawl surveys (DATRAS).” The International Council for the Exploration of the Sea, Copenhagen. **Available at: http://www.emodnet-biology.eu/data-catalog?%3Fmodule=dataset&dasid=2767**, accessed 2013 (2010).
95. DATRAS “Fish trawl survey: Northern Irish Ground Fish Trawl Survey. ICES Database of trawl surveys (DATRAS).” The International Council for the Exploration of the Sea, Copenhagen. **Available at: http://www.emodnet-biology.eu/data-catalog?%3Fmodule=dataset&dasid=2764**, accessed 2013 (2010).
96. DATRAS “Fish trawl survey: Irish Ground Fish Survey for commercial fish species. ICES Database of trawl surveys (DATRAS).” The International Council for the Exploration of the Sea, Copenhagen. **Available at: http://www.emodnet-biology.eu/data-catalog?module=dataset&dasid=2762**, accessed 2013 (2010).

97. DATRAS "Fish trawl survey: ICES French Southern Atlantic Bottom Trawl Survey for commercial fish species. ICES Database of trawl surveys (DATRAS)." The International Council for the Exploration of the Sea, Copenhagen. **Available at: http://www.emodnet-biology.eu/data-catalog?%3Fmodule=dataset&dasid=2759**, accessed 2013 (2010).

98. DATRAS "Fish trawl survey: ICES Beam Trawl Survey for commercial fish species. ICES Database of trawl surveys (DATRAS)." The International Council for the Exploration of the Sea, Copenhagen. **Available at: http://www.emodnet-biology.eu/data-catalog?%3Fmodule=dataset&dasid=2761**, accessed 2013 (2010).

99. DATRAS "Fish trawl survey: ICES North Sea International Bottom Trawl Survey for commercial fish species. ICES Database of trawl surveys (DATRAS)." The International Council for the Exploration of the Sea, Copenhagen. **Available at: http://www.emodnet-biology.eu/data-catalog?%3Fmodule=dataset&dasid=2763**, accessed 2013 (2010).

100. Reichert, M. "MARMAP Fly Net 1990-2009". SCDNR/NOAA MARMAP Program, SCDNR MARMAP Aggregate Data Surveys, The Marine Resources Monitoring, Assessment, and Prediction (MARMAP) Program, Marine Resources Research Institute, South Carolina Department of Natural Resources USA. **Available at: http://www.usgs.gov/obis-usa/**, accessed 2013 (2010).

101. Reichert, M. "MARMAP Yankee Trawl 1990-2009". SCDNR/NOAA MARMAP Program, SCDNR MARMAP Aggregate data surveys, The Marine Resources Monitoring, Assessment, and Prediction (MARMAP) Program, Marine Resources Research Institute, South Carolina Department of Natural Resources USA. **Available at: http://www.usgs.gov/obis-usa**, accessed 2013 (2010).

102. "Northeast Fisheries Science Center Bottom Trawl Survey Data (OBIS-USA)" NOAA's National Marine Fisheries Service (NMFS) Northeast Fisheries Science Center. Woods Hole, Massachusetts, USA. **Available at: http://www.iobis.org/mapper/?dataset=1435**, accessed 2013 (2005).

103. Harmon, M. & Franklin, J. "Long-term growth, mortality and regeneration of trees in permanent vegetation plots in the Pacific Northwest, 1910 to present." Long-Term Ecological Research. Forest Science Data Bank, Corvallis. **Available at: http://andrewsforest.oregonstate.edu/data/abstract.cfm?dbcode=TV010**, accessed 2012 (2012).

104. HMANA "Hawk Migration Association of North America (HMANA)." **Available at: http://www.hmana.org/**, accessed 2012.

105. NatureCounts "Ontario Breeding Bird Atlas (2001-2005): point count data." NatureCounts, a node of the Avian Knowledge Network. Bird Studies Canada. **Available at: http://www.birdscanada.org/birdmon/**, accessed 2012.

106. USFS "Landbird Monitoring Program (UMT-LBMP)." US Forest Service. **Available at: http://www.avianknowledge.net/**, accessed 2012.

107. NatureCounts "Maritimes Breeding Bird Atlas (2006-2010): point count data." NatureCounts, a node of the Avian Knowledge Network. Bird Studies Canada. **Available at: http://www.birdscanada.org/birdmon/**, accessed 2012.

108. Bird Studies Canada "Marsh Monitoring Program - Amphibian Surveys." NatureCounts, a node of the Avian Knowledge Network. **Available at: http://www.birdscanada.org/birdmon/**, accessed 2012 (2012).

109. Bird Studies Canada "Marsh Monitoring Program - Bird Survey." NatureCounts, a node of the Avian Knowledge Network. **Available at: http://www.birdscanada.org/birdmon/**, accessed 2012 (2012).

110. Viereck, L. A., Van Cleve, K., Chapin, F. S., Ruess, R. W. & Hollingsworth, T. N. Vegetation Plots of the Bonanza Creek LTER Control Plots: Species Count (1975 - 2004). Environmental Data Initiative. **Available at: http://dx.doi.org/10.6073/pasta/8dd0e1ac48e2f82b51adabfbd3c62ae2**, accessed 2012 (2005).

111. Shochat, E., Katti, M. & Warren, P. "Point count bird censusing: long-term monitoring of bird distribution and diversity in central Arizona-Phoenix: period 2000 to 2011". Central Arizona-Phoenix Long-Term Ecological Research. Global Institute for Sustainability, Arizona State University. **Available at: https://caplter.asu.edu/data/data-catalog/?id=46**, accessed 2012 (2004).

112. Trexler, J. "Consumer Stocks: Fish, Vegetation, and other Non-physical Data from Everglades National Park (FCE), South Florida from February 2000 to Present." Florida Coastal Everglades LTER Program. **Available at: http://fcelter.fiu.edu/data/core/metadata/EML/?datasetid=LT_CD_Trexler_001**, accessed 2012 (2007).

113. Williams, D. "Pelagic Fish Observations 1968-1999." Australian Antarctic Data Centre. **Available at: http://www.gbif.org/dataset/85b0a82a-f762-11e1-a439-00145eb45e9a**, accessed 2012.

114. Battles, J. J., Fahey, T. & Cleavitt, N. "Forest Inventory of a Northern Hardwood Forest: Watershed 6 1982, Hubbard Brook Experimental Forest." The Hubbard Brook Ecosystem Study LTER Program. **Available at: http://www.hubbardbrook.org/data/dataset.php?id=31**, accessed 2016.

115. Battles, J. J., Fahey, T. & Cleavitt, N. "Forest Inventory of a Northern Hardwood Forest: Watershed 6 1965, Hubbard Brook Experimental Forest." The

Hubbard Brook Ecosystem Study LTER Program. **Available at: http://www.hubbardbrook.org/data/dataset.php?id=29**, accessed 2016.

116. Battles, J. J., Fahey, T. & Cleavitt, N. "Forest Inventory of a Northern Hardwood Forest: Watershed 6 1977, Hubbard Brook Experimental Forest." The Hubbard Brook Ecosystem Study LTER Program. **Available at: http://www.hubbardbrook.org/data/dataset.php?id=30**, accessed 2016.

117. Battles, J. J., Fahey, T. & Cleavitt, N. "Forest Inventory of a Northern Hardwood Forest: Watershed 6 1987, Hubbard Brook Experimental Forest." The Hubbard Brook Ecosystem Study LTER Program. **Available at: http://www.hubbardbrook.org/data/dataset.php?id=32**, accessed 2016.

118. Battles, J. J., Fahey, T. & Cleavitt, N. "Forest Inventory of a Northern Hardwood Forest: Watershed 6 1992, Hubbard Brook Experimental Forest." The Hubbard Brook Ecosystem Study LTER Program. **Available at: http://www.hubbardbrook.org/data/dataset.php?id=33**, accessed 2016.

119. Battles, J. J., Fahey, T. & Cleavitt, N. "Forest Inventory of a Northern Hardwood Forest: Watershed 6 1997, Hubbard Brook Experimental Forest." The Hubbard Brook Ecosystem Study LTER Program. **Available at: http://www.hubbardbrook.org/data/dataset.php?id=34**, accessed 2016.

120. Battles, J. J., Johnson, C., Hamburg, S., Fahey, T., Driscoll, C. & Likens, G. (2003) "Forest Inventory of a Northern Hardwood Forest: Watershed 6 2002." The Hubbard Brook Ecosystem Study LTER Program. **Available at: http://www.hubbardbrook.org/data/dataset.php?id=35**, accessed 2012.

121. Muldavin, E. "Pinon-Juniper (Core Site) Quadrat Data for the Net Primary Production Study at the Sevilleta National Wildlife Refuge, New Mexico (2003-

Present).” Sevilleta Long Term Ecological Research Program. **Available at: http://sev.lternet.edu/node/1718**, accessed 2013.

122. Paquette, A., Laliberté, E., Bouchard, A., Blois, S. de, Legendre, P. & Brisson, J. Lac Croche understory vegetation data set (1998–2006). *Ecology*, **88**, 3209 (2007).

123. Day, F. “Long-term N-fertilized vegetation plots on Hog Island, Virginia Coastal Barrier Islands, 1992-2014.” Virginia Coast Reserve Long-Term Ecological Research Project. **Available at: http://www.vcrlter.virginia.edu/cgi-bin/showDataset.cgi?docid=knb-lter-vcr.106**, accessed 2013 (2010).

124. Day, F. P., Conn, C., Crawford, E. & Stevenson, M. Long-term effects of nitrogen fertilization on plant community structure on a coastal barrier island dune chronosequence. *Journal of Coastal Research*, **20**, 722–730 (2004).

125. Bird Studies Canada “BC Coastal Waterbird Survey (2004).” NatureCounts, a node of the Avian Knowledge Network. **Available at: http://www.birdscanada.org/birdmon/**, accessed 2012 (2012).

126. Chen, H., Liao, Y.-C., Chen, C.-Y., Tsai, J.-I., Chen, L.-S. & Shao, K.-T. Long-term monitoring dataset of fish assemblages impinged at nuclear power plants in northern Taiwan. *Scientific data*, **2**, 150071 (2015).

127. Thomsen, P. F., Jørgensen, P. S., Bruun, H. H., Pedersen, J., Riis-Nielsen, T., Jonko, K., Słowińska, I., Rahbek, C. & Karsholt, O. Resource specialists lead local insect community turnover associated with temperature – analysis of an 18-year full-seasonal record of moths and beetles. *Journal of Animal Ecology*, **85**, 251-261 (2016).

128. Reichert, M. “MARMAP Blackfish Trap Survey 1990-2009”. SCDNR/NOAA MARMAP Program. SCDNR MARMAP Aggregate Data Surveys. The Marine Resources Monitoring Assessment and Prediction (MARMAP) Program. Marine

Resources Research Institute. South Carolina Department of Natural Resources USA. **Available at: http://www.usgs.gov/obis-usa/**, accessed 2013 (2010).

129. Woods, K. D. Multi-decade, spatially explicit population studies of canopy dynamics in Michigan old-growth forests. *Ecology*, **90**, 3587 (2009).

130. Reed, D. C. “SBC LTER: Reef: Kelp forest community dynamics: Abundance and size of giant kelp (Macrocystis pyrifera), ongoing since 2000”. Santa Barbara Coastal LTER. **Available at: http://sbc.lternet.edu/cgi-bin/showDataset.cgi?docid=knb-lter-sbc.18**, accessed 2016 (2014a).

131. Reed, D. C. “SBC LTER: Reef: Kelp forest community dynamics: Fish abundance”. Santa Barbara Coastal LTER. **Available at: http://sbc.lternet.edu/cgi-bin/showDataset.cgi?docid=knb-lter-sbc.17**, accessed 2016 (2014b).

132. Reed, D. C. “SBC LTER: Reef: Kelp forest community dynamics: Invertebrate and algal density”. Santa Barbara Coastal LTER. **Available at: http://sbc.lternet.edu/cgi-bin/showDataset.cgi?docid=knb-lter-sbc.19**, accessed 2016 (2014c).

133. Davis, R. A. & Doherty, T. S. Rapid Recovery of an Urban Remnant Reptile Community following Summer Wildfire. *PLoS ONE*, **10**, e0127925 (2015).

134. Bonebrake, T. C., Pickett, E. J., Tsang, T. P., Tak, C. Y., Vu, M. Q. & Van Vu, L. Warming threat compounds habitat degradation impacts on a tropical butterfly community in Vietnam. *Global Ecology and Conservation*, **8**, 203-211 (2016).

135. Vu, L. V. Diversity and similarity of butterfly communities in five different habitat types at Tam Dao National Park, Vietnam. *Journal of Zoology*, **277**, 15–22 (2009).

136. Edgar, G. J. & Stuart-Smith, R. D. Systematic global assessment of reef fish communities by the Reef Life Survey program. *Nature Scientific Data*, **1**, 140007 (2014).

137. Carpenter, R. "MCR LTER: Coral Reef: Long-term Population and Community Dynamics: Other Benthic Invertebrates, ongoing since 2005". Moorea Coral Reef LTER, knb-lter-mcr.7.28. **Available at: doi:10.6073/pasta/8e7b3a0c7a8bf315739921861cc79d10**, accessed 2016 (2015).

138. Landis, D. & Gage, S. Insect Populations via Sticky Traps at KBS-LTER. **Available at: http://lter.kbs.msu.edu/datatables/67**, accessed 2016 (2014).

139. Joern, A. CGR02 Sweep Sampling of Grasshoppers on Konza Prairie LTER watersheds (1982-present). Environmental Data Initiative. **Available at: http://dx.doi.org/10.6073/pasta/7060b2c244229a37e3bfc8c18f14ad02**, accessed 2016 (2016).

140. Jonas, J. L. & Joern, A. Grasshopper (Orthoptera: Acrididae) communities respond to fire, bison grazing and weather in North American tallgrass prairie: a long-term study. *Oecologia*, **153**, 699-711 (2007).

141. Wiley, R. H. "Population estimates of Appalachian salamanders". Coweeta LTER. **Available at: http://coweeta.uga.edu/eml/1044.xml**, accessed 2016.

142. Merritt, J. Long Term Mammal Data from Powdermill Biological Station 1979-1999. Environmental Data Initiative. **Available at: http://dx.doi.org/10.6073/pasta/83c888854e239a79597999895bb61cfe**, accessed 2016 (1999).

143. Pollard, E., Hall, M. L & Bibby, T. J. Monitoring the Abundance of Butterflies 1976-1985. Research & survey in nature conservation. **Available at: http://jncc.defra.gov.uk/page-2614**, accessed 2016 (1986).

144. Sal, S., López-Urrutia, Á., Irigoien, X., Harbour, D. S. & Harris, R. P. Marine microplankton diversity database. *Ecology*, **9**4, 1658 (2013).

145. Kaufman, D. W. Seasonal summary of numbers of small mammals on 14 LTER traplines in prairie habitats at Konza Prairie. Konza Prairie Long-Term Ecological Research. **Available at: http://lter.konza.ksu.edu/content/csm01-seasonal-summary-numbers-small-mammals-14-lter-traplines-prairie-habitats-konza**, accessed 2016.

146. Prins, H. H. T. & Douglas-Hamilton, I. Stability in a Multi-Species Assemblage of Large Herbivores in East Africa. *Oecologia*, **83**, 392–400 (1990).

147. Knops, J. & Tilman, D. Successional Dynamics on a Resampled Chronosequence - Experiment 014. Cedar Creek Ecosystem Science Reserve. **Available at http://www.cedarcreek.umn.edu/research/data/dataset?ghe014**, accessed 2016.

148. Lightfoot, D. "Lizard pitfall trap data (LTER-II, LTER-III)". Jornada Basin LTER. **Available at: http://jornada.nmsu.edu/lter/dataset/49821/view**, accessed 2016 (2013).

149. Twilley, R., Rivera-Monroy, V. H. & Castaneda, E. "Mangrove Forest Growth from the Shark River Slough, Everglades National Park (FCE), South Florida from January 1995 to Present". Florida Coastal Everglades LTER. **Available at: http://fcelter.fiu.edu/data/core/metadata/?datasetid=LT_PP_Rivera_002**, accessed 2016 (2005).

150. SANParks "Karoo National Park Census Data. 1994 - 2009". **Available at: http://datadryad.org/handle/10255/dryad.13079?show=full**, accessed 2016 (2011).

151. Wilgers, D. J., Horne, E. A., Sandercock, B. K. & Volkmann, A. W. Effects of rangeland management on community dynamics of the herpetofauna of the tallgrass prairie. *Herpetologica*, **62**, 378–388 (2006).

152. Lightfoot, D. & Schooley, R. L. "SMES rodent trapping data, Small Mammal Exclosure Study". Jornada LTER. **Available at: http://jornada.nmsu.edu/sites/jornada.nmsu.edu/files/data_files/JornadaStudy_086_smes_rodent_trapping_data_0.csv**, accessed 2016.

153. Venturoli, F., Felfili, J. M. & Fagg, C. W. Temporal evaluation of natural regeneration in a semideciduous secondary forest in Pirenopolis, Goias, Brazil. *Revista Arvore*, **35**, 473–483 (2011).

154. Kelt, D. A., Meserve, P. L., Gutiérrez, J. R., Milstead, W. B. & Previtali, M. A. Long-term monitoring of mammals in the face of biotic and abiotic influences at a semiarid site in north-central Chile. *Ecology*, **94**, 977 (2013).

155. Pélissier, R., Pascal, J.-P., Ayyappan, N., Ramesh, B. R., Aravajy, S. & Ramalingam, S. R. Tree demography in an undisturbed Dipterocarp permanent sample plot at Uppangala, Western Ghats of India. *Ecology*, **92**, 1376 (2011).

156. Svensson, S., Thorner, A. M. & Nyholm, N. E. I. Species trends, turnover and composition of a woodland bird community in southern Sweden during a period of 57 years. *Ornis Svecica*, **20**, 31–44 (2010).

157. Lightfoot, D. "Small Mammal Exclosure Study (SMES) Vegetation Data from the Chihuahuan Desert Grassland and Shrubland at the Sevilleta National Wildlife Refuge, New Mexico (2006-2009)". Long Term Ecological Research Network. **Available at: http://dx.doi.org/10.6073/pasta/d80d5e2196cd11ef79df23ebe5a77c19**, accessed 2016 (2011).

158. Carvalho, F., Zocche, J. J. & Mendonça, R. Á. Morcegos (Mammalia, Chiroptera) em restinga no município de Jaguaruna, sul de Santa Catarina, Brasil. *Biotemas*, **22**, 193-201 (2009).

159. Davies, C. H., Coughlan, A., Hallegraeff, G., Ajani, P., Armbrecht, L., Atkins, N., Bonham, P., Brett, S., Brinkman, R., Burford, M., Clementson, L., Coad, P., Coman, F., Davies, D., Dela-Cruz, J., Devlin, M., Edgar, S., Eriksen, R., Furnas, M., Hassler, C., Hill, D., Holmes, M., Ingleton, T., Jameson, I., Leterme, S. C., Lønborg, C., McLaughlin, J., McEnnulty, F., McKinnon, A. D., Miller, M., Murray, S., Nayar, S., Patten, R., Pritchard, T., Proctor, R., Purcell-Meyerink, D., Raes, E., Rissik, D., Ruszczyk, J., Slotwinski, A., Swadling, K. M., Tattersall, K., Thompson, P., Thomson, P., Tonks, M., Trull, T.W., Uribe-Palomino, J., Waite, A. M., Yauwenas, R., Zammit, A. & Richardson, A. J. A database of marine phytoplankton abundance, biomass and species composition in Australian waters. *Scientific Data*, **3**, 160043 (2016).

160. Bradford, M. G., Murphy, H. T., Ford, A. J., Hogan, D. L. & Metcalfe, D. J. Long-term stem inventory data from tropical rain forest plots in Australia. *Ecology*, **95**, 2362-2362 (2014).

161. Stapp, P. SGS-LTER Long-Term Monitoring Project: Small Mammals on Trapping Webs on the Central Plains Experimental Range, Nunn, Colorado, USA 1994 -2006, ARS Study Number 118. Environmental Data Initiative. **Available at: http://dx.doi.org/10.6073/pasta/2e311b4e40fea38e573890f473807ba9**, accessed 2017 (2013).

162. Dickson, J. G., Conner, R. N. & Williamson, J. H. Neotropical migratory bird communities in a developing pine plantation. Proceedings on the Annual Conference. *SEAFWA*, **47**, 439-446 (1993).

163. Reed, D. C. “SBC LTER: Reef: Kelp Forest Community Dynamics: Fish abundance”. Santa Barbara Coastal LTER. **Available at: doi:10.6073/pasta/e37ed29111b2fddffc08355252b8b8c7**, accessed 2016 (2014).

164. Hall, G. A. A Long-Term Bird Population Study in an Appalachian Spruce Forest. *The Wilson Bulletin*, **96**, 228–240 (1984).

165. Enemar, A., Sjöstrand, B. E., Andersson, G. Ö. & von Proschwitz, T. The 37-year dynamics of a subalpine passerine bird community, with special emphasis on the influence of environmental temperature and Epirrita autumnata cycles. *Ornis Svecica*, **14**, 63–106 (2004).

166. Willis, T. “Hahei marine dataset (1997-2002), New Zealand fish”. Institute of Marine Sciences, University of Portsmouth. Accessed 2016.

167. Lightfoot, D. “Small Mammal Exclosure Study (SMES)”. Sevilleta Long Term Ecological Research Program. **Available at: http://sev.lternet.edu/content/small-mammal-exclosure-study-smes-0**, accessed 2016.

168. Monitoring Site 1000 Project, Biodiversity Center, Ministry of Environment of Japan (2015) “Monitoring site 1000 Coastal zone research - Tidal flat survey” (**HIG01.zip, downloaded from http://www.biodic.go.jp/moni1000/findings/data/index_file_tidalflats.html**). Accessed 2016.

169. Monitoring Site 1000 Project, Biodiversity Center, Ministry of Environment of Japan (2015) “Monitoring site 1000 Alpine research - Surface wandering beetles” (**KOZ07zip, downloaded from http://www.biodic.go.jp/moni1000/findings/data/index.html**). Accessed 2016.

170. Monitoring Site 1000 Project, Biodiversity Center, Ministry of Environment of Japan (2014) “Monitoring site 1000 Village survey - Bird survey data (2005-2012)”

(**SAT02.zip, downloaded from http://www.biodic.go.jp/moni1000/findings/data/index.html**). Accessed 2016.

171. Monitoring Site 1000 Project, Biodiversity Center, Ministry of Environment of Japan (2014) “Monitoring site 1000 Village survey - Medium and large mammal survey data (2006-2012)” (**SAT03zip, downloaded from http://www.biodic.go.jp/moni1000/findings/data/index.html**). Accessed 2016.

172. Monitoring Site 1000 Project, Biodiversity Center, Ministry of Environment of Japan (2013) “Monitoring site 1000 Shorebird Survey” (**ShorebirdsDatapackage2012.zip, downloaded from http://www.biodic.go.jp/moni1000/ findings/data/index.html**). Accessed 2016.

173. Monitoring Site 1000 Project, Biodiversity Center, Ministry of Environment of Japan (2014) “Monitoring site 1000 Forest and grassland research - Surface wandering beetles survey data” (**GBDataPackage2014ver1.zip, downloaded from http://www.biodic.go.jp/moni1000/findings/data/index.html**). Accessed 2016.

174. Monitoring Site 1000 Project, Biodiversity Center, Ministry of Environment of Japan (2014) “Monitoring site 1000 Forest and grassland research - Surface wandering beetles survey data” (**GBDataPackage2014ver1.zip, downloaded from http://www.biodic.go.jp/moni1000/findings/data/index.html**). Accessed 2016.

175. Benedetti-Cecchi, L. “Calafuria Mid-shore Intertidal Dataset (1991-2014)”. Department of Biology, University of Pisa. Accessed 2016.

176. Benedetti-Cecchi, L. “Calafuria Low-shore Intertidal Dataset (1991-2014)”. Department of Biology, University of Pisa. Accessed 2016.

177. NERC “The Global Population Dynamics Database Version 2”. Centre for Population Biology, Imperial College. **Available at: http://www.sw.ic.ac.uk/cpb/cpb/gpdd.html**, accessed 2016 (2010).

178. Pollard, E. Monitoring butterfly numbers. In: F. B.Goldsmith (ed) Monitoring for Conservation and Ecology. Chapman and Hall (1991).

179. How, R. A. Long-term sampling of a herpetofaunal assemblage on an isolated urban bushland remnant, Bold Park, Perth. *Journal of the Royal Society of Western Australia*, **81**, 143-148 (1998).

180. Krefting, L. W. & Ahlgren, C. E. Small Mammals and Vegetation Changes After Fire in a Mixed Conifer-Hardwood Forest. *Ecology*, **65**, 1391–1398 (1974).

181. Hsieh, C.-H. "Icthyoplankton data collected from Yenliao Bay in 6 stations northeast of Taiwan (1995-2000)". Ecoinformatics Lab, Institute of Oceanography National Taiwan University. Accessed 2016.

182. Kendeigh, S. C. (1982) Bird populations in east central Illinois: Fluctuations, variations, and development over a half-century. University of Illinois Press.

183. Fraser, W. "At-sea seabird censuses. Data on the species encountered (including marine mammals), their abundance, distribution and behavior. Data collected aboard cruises off the coast of the Western Antarctic Peninsula, 1993 - present". Palmer Station Antarctica LTER. **Available at: http://dx.doi.org/10.6073/pasta/e3871e749fa737dd94d5a269ac90e8ce**, accessed 2016 (2014).

184. Svensson, S. Species composition and population fluctuations of alpine bird communities during 38 years in the Scandinavian mountain range. *Ornis Svecica*, **16**, 183–210 (2006).

185. Monitoring Site 1000 Project, Biodiversity Center, Ministry of Environment of Japan (2015) "Monitoring site 1000 Alpine research - Butterfly Survey" (**KOZ06.zip, downloaded from http://www.biodic.go.jp/moni1000/findings/data/index.html**). Accessed 2016.

186. Monitoring Site 1000 Project, Biodiversity Center, Ministry of Environment of Japan (2015) “Monitoring site 1000 Alpine research - Bumblebee Survey” (**KOZ08.zip, downloaded from http://www.biodic.go.jp/moni1000/findings/data/index.html**). Accessed 2016.

187. Barceló, C., Ciannelli, L., Olsen, E. M., Johannessen, T. & Knutsen, H. Eight decades of sampling reveal a contemporary novel fish assemblage in coastal nursery habitats. *Global change biology*, **22**, 1155-1167 (2016).

188. Olsen, E. M., Carlson, S. M., Gjøsæter, J. & Stenseth, N. C. Nine decades of decreasing phenotypic variability in Atlantic cod. *Ecology Letters*, **12**, 622–631 (2009).

189. Rogers, L. A., Stige, L. C., Olsen, E. M., Knutsen, H., Chan, K.-S. & Stenseth, N. C. Climate and population density drive changes in cod body size throughout a century on the Norwegian coast. *Proceedings of the National Academy of Sciences*, **108**, 1961–1966 (2011).

190. Stenseth, N. C., Bjørnstadf, O. N., Falck, W, Fromentin, J. M., Gjøsieter, J. & Gray, J. S. Dynamics of coastal cod populations: intra-and intercohort density dependence and stochastic processes. *Proceedings of the Royal Society of London B: Biological Sciences*, **266**, 1645–1654 (1999).

191. Steinberg, D. Zooplankton collected with a 2-m, 700-um net towed from surface to 120 m, aboard Palmer Station Antarctica LTER annual cruises off the western antarctic peninsula, 2009 - 2016. Environmental Data Initiative. **Available at: http://dx.doi.org/10.6073/pasta/fb658789188724be5f27c81a634647d5**, accessed 2017 (2017).

192. Hoey, A. “Karimunjawa WCS fish data”. Accessed 2016.

193. Hoey, A. “Aceh WCS fish surveys”. Accessed 2016.

194. Zakharov, V. D. Biodiversity of bird population of terrestrial habitats in Southern Ural. Miass: IGZ, Ural Branch of Russian Academy of Sciences, 158 p (1998).

195. Berezovikov, N. N. The birds of settlements in Markakol Depression (Southern Altai). *Russian Ornithological Journal,* **249**, 3-15 (2004).

196. Melnikov, Y. I., Melnikova, N. & Pronkevich, V. V. Migration of birds of prey in the mouth of the river Irkut. *Russian Ornithological Journal*, **108**, 3–17 (2000).

197. Nedosekin, V. Y. Long-term dynamics of the population and the quantity of small mammals under conditions of the reserve "Galichya Gora". *Proceedings of National Nature Reserve Prisursky*, **30**, 87–90 (2015).

198. Thorn, S., Bässler, C., Bernhardt-Römermann, M., Cadotte, M., Heibl, C., Schäfer, H., Seibold, S. & Müller, J. Changes in the dominant assembly mechanism drive species loss caused by declining resources. *Ecology Letters*, **19**, 163–170 (2016).

199. Thorn, S., Bässler, C., Gottschalk, T., Hothorn, T., Bussler, H., Raffa, K. & Müller, J. New insights into the consequences of post-windthrow salvage logging revealed by functional structure of saproxylic beetles assemblages. *PLoS ONE,* **9**, e101757 (2014).

200. Thorn, S., Werner, S. A., Wohlfahrt, J., Bässler, C., Seibold, S., Quillfeldt, P. & Müller, J. Response of bird assemblages to windstorm and salvage logging - Insights from analyses of functional guild and indicator species. *Ecological Indicators*, **65**, 142–148 (2016).

201. Neat, F. & Campbell, N. Demersal fish diversity of the isolated Rockall plateau compared with the adjacent west coast shelf of Scotland. *Biological Journal of the Linnean Society*, **104**, 138–147 (2011).

202. Kenner, M. C., Estes, J. A., Tinker, M. T., Bodkin, J. L., Cowen, R. K., Harrold, C., Hatfield, B. B., Novak, M., Rassweiler, A. & Reed, D. C. A multi-decade time series of kelp forest community structure at San Nicolas Island, California (USA). *Ecology*, **94**, 2654 (2013).

203. Kushner, D. J., Rassweiler, A., McLaughlin, J. P. & Lafferty, K. D. A multi-decade time series of kelp forest community structure at the California Channel Islands. *Ecology*, **94**, 2655 (2013).

204. Muldavin, E. & Collins, S. Prescribed Burn Effect on Chihuahuan Desert Grasses and Shrubs at the Sevilleta National Wildlife Refuge, New Mexico: Species Composition Study 2004 to present. Sevilleta LTER. **Available at: http://sev.lternet.edu/data/sev-166**, accessed 2016 (2003).

205. Anderson, J., Vermeire, L. & Adler, P. B. Fourteen years of mapped, permanent quadrats in a northern mixed prairie, USA. *Ecology*, **92**, 1703-1703 (2011).

206. Hogstad, O. Structure and dynamics of a passerine bird community in a spruce-dominated boreal forest. A 12-year study. *Annales Zoologici Fennici*, **30**, 43-54 (1993).

207. Douglass, J. G., France, K. E., Richardson, J. P. & Duffy, J. E. Seasonal and interannual change in a Chesapeake Bay eelgrass community: Insights into biotic and abiotic control of community structure. *Limnology and Oceanography*, **55**, 1499–1520 (2010).

208. Lefcheck, J. S. The use of functional traits to elucidate the causes and consequences of biological diversity. The College of William & Mary, PhD thesis. **Available at: http://gradworks.umi.com/36/62/3662989.html**, accessed 2016 (2015).

209. Institute of Agricultural and Fisheries research (ILVO), Belgium Macrobenthos monitoring at long-term monitoring stations in the Belgian part of the North Sea between 1979 and 1999. **Available at: http://dx.doi.org/10.14284/201**, accessed 2016 (2016).

210. Institute of Agricultural and Fisheries research (ILVO), Belgium Macrobenthos monitoring at long-term monitoring stations in the Belgian part of the North Sea from 2001 on. **Available at: http://dx.doi.org/10.14284/202**, accessed 2016 (2016).

211. Woods, K. D. Multi-decade biomass dynamics in an old-growth hemlock-northern hardwood forest, Michigan, USA. *PeerJ*, **2**, e598 (2014).

212. Belmaker, J., Ziv, Y. & Shashar, N. The influence of connectivity on richness and temporal variation of reef fishes. *Landscape ecology*, **26**, 587-597 (2011).

213. Edelist, D., Rilov, G., Golani, D., Carlton, J. T. & Spanier, E. Restructuring the Sea: profound shifts in the world's most invaded marine ecosystem. *Diversity and Distributions*, **19**, 69–77 (2013).
